# Coxsackievirus B Escapes Antiviral CD8⁺ T Cells but Triggers Robust CD4⁺ Memory Responses

**DOI:** 10.64898/2026.01.16.699852

**Authors:** Orlando Burgos-Morales, Federica Vecchio, Fatoumata Samassa, Margot Petit, Zhicheng Zhou, Barbara Brandao, Alexia Carré, Annalisa Nicastri, Valeriia Dotsenko, Chloe Shepherd, Keijo Viiri, Robert Parker, Amelia K. Linnemann, Sylvaine You, Malin Flodstrom-Tullberg, Nicola Ternette, Roberto Mallone

## Abstract

Coxsackieviruses B (CVBs) are plausible triggers of the pancreatic islet autoimmunity leading to type 1 diabetes. Islet autoantibody seroconversion correlates with persistent CVB infections in the gut and pancreas, suggesting defective antiviral control and the need to define immune mechanisms at the intestinal entry site. We investigated how CVB3 modulates antigen presentation, the viral immunopeptidome of enterocytes and antigen-presenting cells, and downstream T-cell immunity. CVB3-infected enterocytes escaped immune recognition by downregulating HLA Class I and viral peptide presentation, impairing CD8^+^ T-cell responses in vitro. In CVB-seropositive individuals, circulating CVB-reactive CD8^+^ T cells were stalled in naïve-like and exhausted effector/memory states. In contrast, CVB3 induced HLA class II upregulation, promoting robust CD4⁺ T-cell activation. Circulating CVB3-reactive CD4⁺ T cells fully differentiated into polyfunctional T helper memory. These findings indicate that CVB3 antiviral control is predominantly CD4⁺ T-cell-mediated and provide a rationale for mucosal vaccination strategies and immune monitoring tools to follow infection or vaccination.

**Teaser:** Coxsackievirus evades CD8+ T-cell immunity in the gut, leaving CD4+ T cells as the main line of antiviral defense.

## Introduction

Type 1 diabetes (T1D) is an autoimmune disease of unknown etiology. Its increasing incidence, including in individuals harboring neutral and protective human leukocyte antigen Class II (HLA-II) haplotypes, suggests that environmental triggers may play a primary role in disease onset (*1*). These triggers likely have their greatest impact early in life, as a large proportion of children who later develop T1D exhibit islet autoantibody (aAb) seroconversion before the age of 2 (*2*).

Histopathological and epidemiological studies point to Coxsackieviruses B (CVBs) as plausible triggers of the autoimmune T-cell attack against pancreatic islet β cells (*3*). The histopathological co-localization of the enteroviral viral protein (VP)1 and double-stranded (ds)RNA with HLA Class I (HLA-I) hyperexpression (a recognized hallmark of disease) in the islets of T1D donors (*4–6*) suggest that low-grade, persistent CVB infections may trigger the autoimmune process (*7*). In line with these observations, stool metagenome sequencing on prospective cohorts of genetically at-risk children demonstrated a significant correlation between prolonged CVB infections (characterized by the persistent shedding of the same CVB serotype in sequential stool samples) and the aAb seroconversion marking the initiation of islet autoimmunity (*8*). These persistent CVB infections (*7*), possibly facilitated by inefficient anti-viral immune responses in predisposed individuals (*3, 9*), may contribute to T1D autoimmunity.

While a causal link between CVBs and the onset of T1D autoimmunity remains elusive, the positive results of recent clinical trials may provide this missing link and novel intervention and prevention strategies by enhancing antiviral responses. The antiviral agents pleconaril and ribavirin provided some preservation of residual insulin secretion in children and adolescents with new-onset clinical (stage 3) T1D (*10*). In addition, T1D primary prevention trials are being considered to explore the protective effect of a multivalent inactivated CVB vaccine (*11–13*). These trials aim to directly assess whether preventing CVB infection by vaccination can offer protection against islet autoimmunity and subsequent T1D development, potentially providing proof for the viral etiology of the disease and a primary prevention option not available to date.

Several knowledge gaps however exist about the natural immune response against CVB (*3*). One serological study suggests that children who later develop anti-insulin aAbs at an early age do not harbor anti-CVB neutralizing antibodies (Abs) (*14*). This could potentially make them more susceptible to infections with higher viral loads, increasing the likelihood of high-load viremia and CVB spreading to the pancreas (*3*). On the same lines, we have previously documented that CVB infects and destroys β cells, but elicits limited antiviral CD8^+^ T-cell responses (*9*). Overall, these observations provide a strong rationale for CVB vaccination.

Since CVB enters the body mainly through gut epithelia before infecting β cells, both tissue types are relevant to understand the mechanisms by which CVB may trigger T1D. Available studies also show that persistent infections in mucosal tissues are strongly associated with T1D (*15, 16*), suggesting that impaired antiviral immunity at the entry site may contribute to disease onset. Yet, the nature of T-cell responses, and how CVB modulates antigen presentation in the intestinal mucosa, remain poorly understood. Here, we addressed this gap by focusing on antigen presentation as a critical determinant of T-cell-mediated immunity, using two complementary models of infection: intestinal epithelial cells and professional antigen-presenting cells (APCs). Through an integrated proteomics and immunopeptidomics strategy, we uncovered viral modulation of antigen processing pathways in enterocytes, characterized by reduced HLA-I but increased HLA-II expression, and a specific repertoire of CVB peptides presented. These experiments were complemented with a macrophage model to map the HLA-II-restricted viral peptides presented upon uptake of cell debris from CVB-infected cells. These datasets enabled us to define novel HLA-A*02:01- and HLA-DRB1*04:01-restricted CVB epitopes and assess their immunogenicity. In line with the differential effect on HLA-I and HLA-II modulation, our results reveal selective impairment of CD8⁺ T-cell responses alongside robust, polyfunctional CD4⁺ T-cell responses, highlighting CVB-driven immune evasion strategies that favor CD4⁺ T cells.

## Results

### CVB infection of CaCo2 enterocytes reveals immune escape mechanisms and hijacking of host cell machinery

To investigate the early intestinal mucosal phase of CVB infection, we established an in-vitro model using CaCo2 enterocytes, which express both HLA-I (A*02:01, A2 from hereon; B*15:01, C*04:01) and HLA-II molecules (DRB1*04:04, DRB4*04:01, DQA1*03:01, DQB1*03:02, DPA1*01:03, DPB1*04:01). We first titrated multiplicities of infection (MOI) using a prototypic CVB3 strain and determined the percentage of infected CaCo2 enterocytes by flow cytometry measurement of the intracellular capsid VP1 and viral replication intermediate double-stranded (ds)RNA at 6 h post-infection (hpi) (*9*). Infection rates increased with MOI, reaching a maximum of ∼49% infected cells at 300 MOI while preserving cell viability (Fig. 1A, left; representative staining profiles in <u>Fig. S1</u>). We then determined whether the proportion of infected cells could be further increased by extending the infection to 8 and 10 hpi. The percentage of infected VP1^+^dsRNA^+^ cells reached a plateau at ∼70% after 10 hpi, again with minimal impact on cell viability (Fig. 1A, right). To validate this infection protocol, we used a CVB3-GFP strain to track infection kinetics in CaCo2 cells by live-cell fluorescence imaging (*9*). Infection (here identified as GFP signal) exhibited an initial lag phase up until 5 hpi, followed by an exponential increase in GFP^+^ cells that plateaued at ∼10 hpi (Fig. 1B), in line with flow cytometry results. Live-cell imaging further revealed morphological changes in infected enterocytes similar to those we previously observed in pancreatic β cells (*9*): cell-to-cell contact through filopodia-like projections allowed cells to transition from non-infected (GFP^−^) to infected (GFP^+^) state (Fig. 1C), suggesting that CVB3-induced cytoskeletal remodeling contributes to viral transfer.

**Figure 1.**
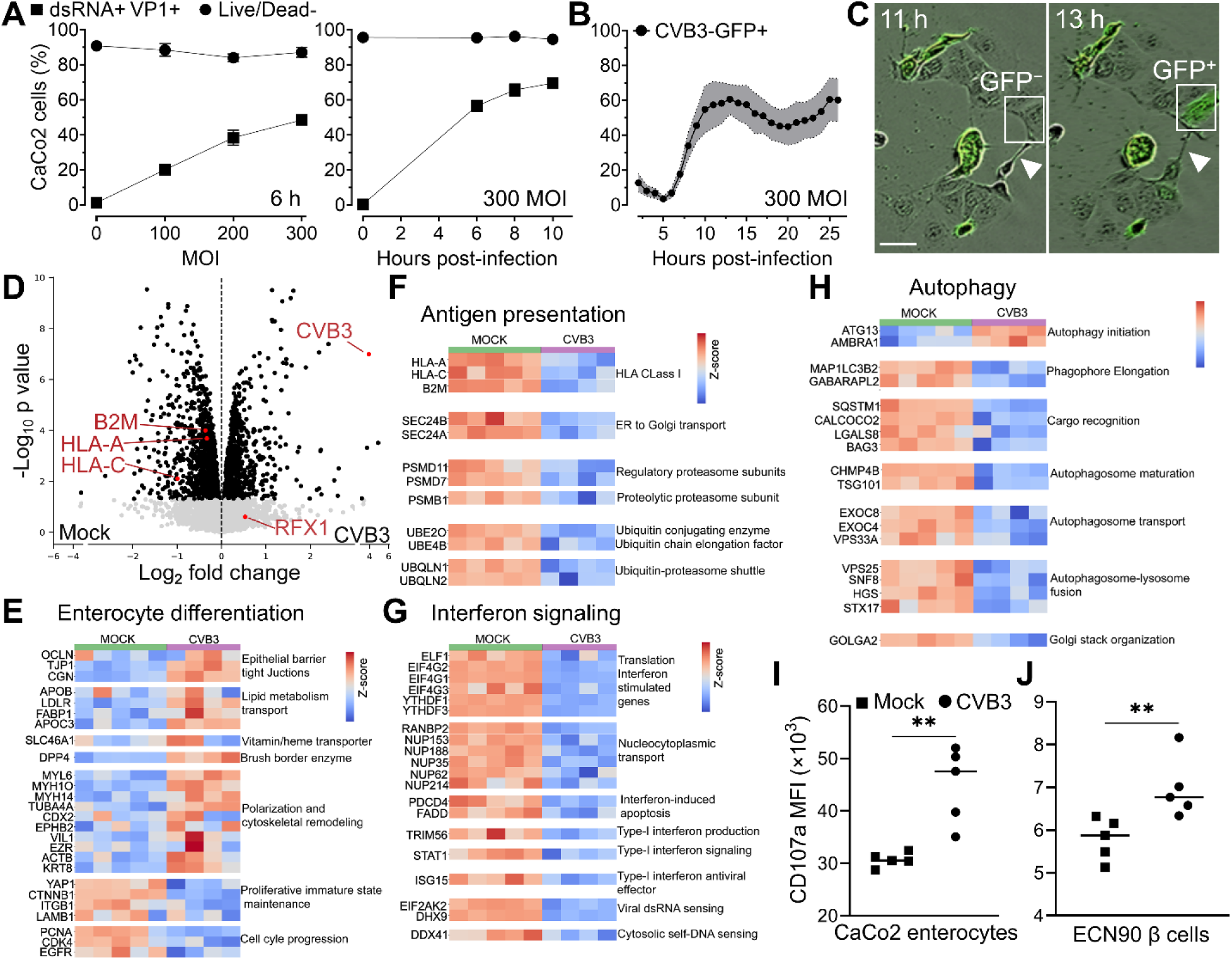
CVB3 immune escape mechanisms in infected CaCo2 enterocytes. **A.** Setup of the in-vitro infection protocol by flow cytometry: optimization of the 300 MOI CVB3 load (*left*; 3 replicates) and 10-h infection time course (*right*; 4 replicates) to achieve maximal infection rates (percent dsRNA^+^VP1^+^ cells) without cell death (percent Live/Dead^−^ cells). **B.** Infection time course using a fluorescent CVB3-GFP strain (300 MOI), measured by live-cell imaging for 25 h (8 replicates). Data in (A-B) represent mean±SD from a representative experiment performed in triplicate. **C.** Phase contrast images overlaid with GFP fluorescence of CVB3-GFP-infected CaCo2 enterocytes at 11 h and 13 h. Arrows point to filopodia. Scale bar 50 µm. **D.** Proteomics analysis of CVB3-infected (n=4) vs. mock-infected (n=5) CaCo2 enterocytes (300 MOI, 10 h). Volcano plot of differentially expressed proteins (black dots) with a significance threshold of p>0.05. **E-H.** Functional categories of proteins differentially expressed: proteins related to enterocyte differentiation (E), antigen processing and presentation (F), IFN signaling (G) and autophagy (H). **I-J.** Surface expression (median fluorescent intensity, MFI) of the lysosomal marker CD107a (LAMP1) in CVB3- vs. mock-infected CaCo2 enterocytes (I; 100 MOI, 20 h) and ECN90 β cells (J; 50 MOI, 16 h). Bars represent median values of 5 replicates from a representative experiment performed in duplicate. **p=0.008 by Mann-Whitney U test.

To understand how CVB3 modulates host cellular pathways, we performed quantitative proteomics on CVB3- vs. mock-infected CaCo2 enterocytes. Among the 6,944 proteins detected, 1,125 (16.2%) were differentially expressed (Fig. 1D). As detailed below, CVB3 infection induced an extensive proteome remodeling, consistent with a coordinated viral strategy to skew enterocyte development and suppress immune recognition.

First, CVB3 infection shifted enterocyte identity towards a more differentiated and polarized phenotype (Fig. 1E). Tight junction proteins (OCLN, TJP1, CGN), cytoskeletal and apical polarity markers (MYH10, MYL6, VIL1, CDX2, TUBA4A, KRT8, ACTB, EZR), and the brush border enzyme DPP4 were upregulated. Lipid and vitamin transport proteins (APOB, LDLR, FABP1, APOC3, SLC46A1) were also increased, suggesting maintenance of absorptive functions despite infection. In contrast, CVB3 infection reduced the expression of key regulators of proliferation and tissue renewal, including YAP1, CTNNB1 (β-catenin), ITGB1 and LAMB1, along with cell cycle effectors PCNA, CDK4 and EGFR. These proteins are involved in maintaining enterocyte progenitor pools (*17*), promoting regenerative signaling (*18*), and driving G1/S cell cycle progression (*19*). This may reflect a viral strategy to arrest the host cell cycle, particularly at the G1 phase, as previously reported in enteroviral infections (*20*). Collectively, these changes maintain a polarized mature phenotype in enterocytes, facilitating viral persistence and altering epithelial barrier integrity due to impaired cell replacement.

Second, multiple components of the antigen processing and presentation pathways were downregulated (Fig. 1F), including HLA-I (HLA-A, HLA-C; but not HLA-B) and β2-microglobulin (B2M), thus impairing surface expression of peptide (p)HLA-I complexes essential for CD8⁺ T-cell recognition. This downregulation extended to the proteasome, with reduced expression of regulatory (PSMD7, PSMD11) and catalytic (PSMB1) subunits needed for antigenic peptide processing. Endoplasmic reticulum (ER)-to-Golgi transport mediators SEC24A and SEC24B, which facilitate HLA-I trafficking, were also reduced. Furthermore, proteins involved in the ubiquitin-proteasome system, specifically the conjugation enzyme UBE2O, elongation factor UBE4B, and shuttling factors UBQLN1/2, were downregulated. These changes highlight how CVB3 disrupts antigen processing and presentation at multiple steps to evade detection by cytotoxic CD8^+^ T cells, echoing similar immune escape features previously observed in infected β cells (*9*).

Third, CVB3 disrupted type I interferon (IFN)-related pathways at multiple levels (Fig. 1G). Host sensors of viral RNA (EIF2AK2, DHX9) and self-DNA (DDX41) were downregulated, indicating impaired recognition of viral replication intermediates and infection-induced cytosolic damage. TRIM56, a key type I IFN-inducing factor, was also reduced, suggesting that CVB3 suppresses IFN production pathways to limit host antiviral responses. Downstream, the IFN signaling axis was disrupted by the loss of STAT1, ISG15 and apoptosis-linked IFN-stimulated genes (ISGs) PDCD4 and FADD, further preventing immune clearance of infected cells. Additionally, several translation regulatory factors (ELF1, EIF4G1/2/3, YTHDF1/3) were downregulated, consistent with a global reduction in host protein synthesis, which may in turn limit ISG expression. Suppression of multiple nucleoporins (RANBP2, NUP153, NUP188, NUP35, NUP62, NUP214) was also evident, potentially disrupting the nuclear trafficking of transcription factors and further attenuating ISG expression. Collectively, these alterations suggest a coordinated viral strategy that disrupts each step of the IFN cascade: from nucleic acid sensing to IFN production and signaling, ultimately impairing ISG translation and facilitating CVB3 replication and spreading under minimal antiviral pressure.

Last, CVB3 infection remodeled autophagy pathways (Fig. 1H). Early autophagy initiators, including ATG13 and AMBRA1, were upregulated, reflecting a host response to the increase of undegraded proteins and organelles caused by inhibited anterograde trafficking (Fig. 1F). This enhanced initiation may increase phagophore formation, providing CVB3 with scaffolds for assembling viral replication organelles (VROs). In parallel, the Golgi structural protein GOLGA2 was downregulated, a change known to drive Golgi disassembly and release of membrane fragments that are often used for VRO assembly (*21–23*). Despite enhanced autophagy initiation, CVB3 infection blocked downstream autophagic progression. Proteins involved in phagophore elongation (MAP1LC3B2, GABARAPL2), cargo recognition (SQSTM1, CALCOCO2, LGALS8, BAG3), autophagosome maturation (CHMP4B, TSG101), vesicle transport (EXOC8, EXOC4, VPS33A), and lysosomal fusion (VPS25, SNF8, HGS, STX17) were all downregulated. This pattern suggests that, while autophagosome formation is triggered, autophagic flux may be arrested, preventing lysosomal degradation of viral components and restricting autophagy-dependent antigen presentation. This inhibition of the autophagy flux often drives cells to use alternative pathways to eliminate autophagosomes, notably by releasing their contents in the extracellular space (*24–26*). Consistent with this, surface cell exposure of the lysosomal marker LAMP1 (CD107a) increased upon CVB3 infection, in CaCo2 enterocytes (Fig. 1I) and also in ECN90 β cells (Fig. 1J).

Collectively, proteome changes in CVB3-infected CaCo2 enterocytes reflect a viral strategy to promote immune silencing and efficient replication. Infected enterocytes adopt a non-proliferative, yet polarized and functionally active, phenotype, concomitantly suppressing antigen processing and presentation, IFN signaling, and autophagy maturation. In addition, filopodia-mediated CVB transmission provides a supplementary mechanism to evade immune surveillance.

### CVB3 infection downregulates HLA-I and upregulates HLA-II antigen presentation in CaCo2 enterocytes

Given the observed proteome downmodulation of antigen presentation and processing pathways in CVB3-infected CaCo2 enterocytes, we next assessed the modulation of surface HLA-I and HLA-II expression by flow cytometry. CVB3 infection induced a sustained dual effect (Fig. 2A): a reduction in HLA-I (∼1.5-fold decrease from 0 hpi) and an induction of HLA-II (∼13-fold increase). Stratifying CaCo2 enterocytes by infection status revealed that these changes were exclusively driven by the VP1^+^ subset, whereas VP1^−^ bystander cells remained unaffected (Fig. 2B). Similarly, significant downregulation of *HLA-A*, *HLA-B*, and *HLA-C* and upregulation of *HLA-DRB1*04* transcripts were observed (Fig. 2C).

**Figure 2.**
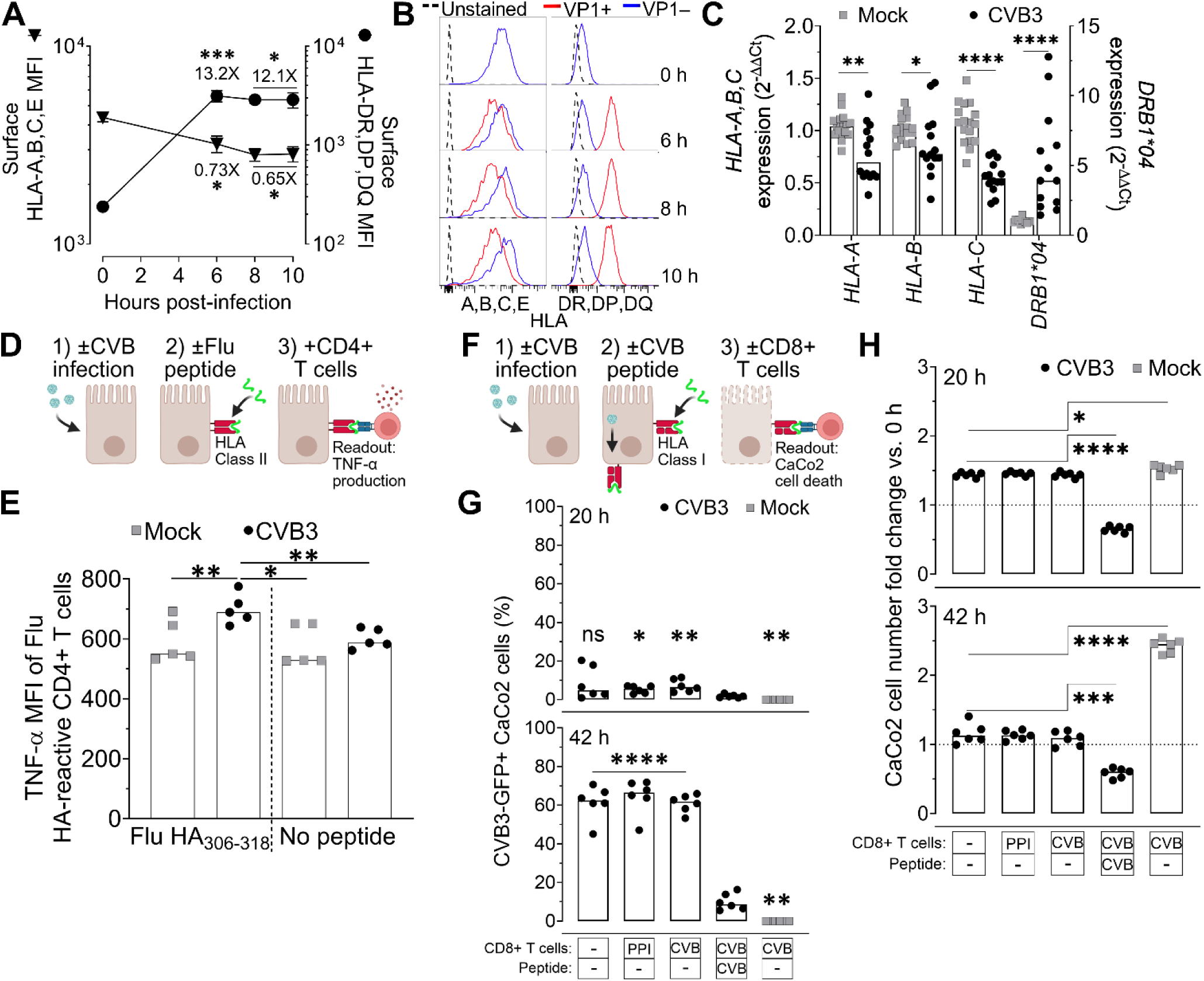
CVB3 infection downregulates HLA-I and upregulates HLA-II antigen presentation in CaCo2 enterocytes. **A.** Surface HLA-I (*left y axis*) and HLA-II expression (*right y axis*) measured by flow cytometry following CVB3 infection (300 MOI) of CaCo2 enterocytes. Data represent mean±SD of 3-4 replicates from a representative experiment performed in triplicate. *p≤0.02, ***p=0.001 vs. 0 h post-infection by Student’s t test. **B.** Representative histograms from the same experiments of surface HLA-I (*left*) and HLA-II (*right*) expression in infected (VP1^+^) vs. non-infected (VP1^−^) CaCo2 enterocytes. **C.** mRNA expression of *HLA-A*, *B*, *C*, and *HLA-DRB1*04* in CVB3- vs. mock-infected CaCo2 enterocytes (300 MOI, 6 h), measured by RT-qPCR and expressed as relative fold change, using *GAPDH* as the housekeeping gene. Bars represent median values of 12 replicates from a representative experiment performed in triplicate. *p=0.005, **p=0.025, ****p≤0.0001 by Mann-Whitney U test. **D.** Schematic of CD4⁺ T-cell activation assays to assess HLA-DR*04:01 upregulation. CVB3- or mock-infected CaCo2 enterocytes (10 MOI, 24 h) were pulsed or not with Flu HA_306-318_ peptide and co-cultured with a Flu HA-reactive CD4⁺ T-cell clone. **E.** CD4^+^ T-cell activation was measured by flow cytometry following intracellular TNF-α staining (expressed as median fluorescence intensity, MFI). Bars represent median values of 5 replicates from a representative experiment performed in triplicate. *p=0.048, **p=0.01 by Student’s t test vs. CVB3-infected and Flu HA peptide-pulsed condition. **F.** Schematic of CD8⁺ T-cell activation assays to assess HLA-A2 downregulation. CVB3-GFP- or mock-infected CaCo2 enterocytes (10 MOI) were pulsed or not with CVB_1246-1254_ peptide and co-cultured or not with irrelevant PPI_15-24_-reactive or cognate CVB-reactive primary CD8^+^ TCR transductants during a 42-h live cell imaging time course to measure infection and CaCo2 cell numbers. **G.** Percent CVB3-GFP^+^ infected CaCo2 enterocytes at 20 h (*top*) and 42 h (*bottom*). *p=0.013, **p≤0.004, ****p≤0.0001 by Student’s t test vs. CVB T cells/CVB peptide condition. **H.** Fold change in the number of live (mKate2^+^) CaCo2 cells (transduced with the mKate2 nuclear marker) normalized to the 0-h input number at 20 h (*top*) and 42 h (*bottom*). *p≤0.014, ***p≤0.0002, ****p≤0.0001 by Student’s t test. Bars in (G-H) represent median values of 6 replicates from a representative experiment performed in duplicate.

We next assessed whether the observed HLA-II upregulation enhanced antigen presentation to CD4^+^ T cells. CaCo2 cells were either CVB3- or mock-infected, pulsed or not with a Flu HA_306-318_ peptide, and co-cultured with a DRB1*04:01-restricted, Flu HA_306-318_-reactive CD4^+^ T-cell clone (Fig. 2D). CVB3-infected, peptide-pulsed CaCo2 cells elicited superior T-cell activation than mock-infected controls, measured by intracellular TNF-α staining (Fig. 2E), demonstrating the functional relevance of HLA-II upregulation.

Similarly, we addressed the functional impact of HLA-I downregulation by measuring CD8^+^ T-cell cytotoxicity by live-cell imaging (Fig. 2F). CaCo2 cells transduced with the nuclear marker mKate2 were infected with CVB3-GFP (10 MOI) and pulsed or not with CVB3_1249-1257_ peptide (KLNSSVYSL; presented by CVB3-infected CaCo2 enterocytes, see Table 1) to enhance the endogenous viral peptide presentation. They were then co-cultured with HLA-A2-restricted T-cell receptor (TCR) transductants (*9*) recognizing either CVB3_1249-1257_ or an irrelevant (preproinsulin, PPI_15-24_) peptide. GFP^+^ cells increased from 0 to 5% at 20 hpi (Fig. 2G, top), marking the onset of viral spreading, and reached 60% at 42 hpi (Fig. 2G, bottom). However, the presence of CVB3-reactive (or PPI-reactive) CD8^+^ T cells did not inhibit viral spreading, unless the peptide was exogenously added, confirming the efficient immune escape afforded by HLA-I downregulation. Accordingly, upon CVB3 infection CaCo2 cell numbers remained stable across conditions irrespective of T-cell co-culture in the absence of CVB3 peptide pulsing (Fig. 2H). At 42 hpi, CVB3-infected CaCo2 cell numbers decreased compared to the mock-infected condition, which underwent some proliferation. This is in line with the reduced expression of proliferation/regeneration-associated proteins upon CVB3 infection observed by proteomics (Fig. 1E).

**Table 1.**
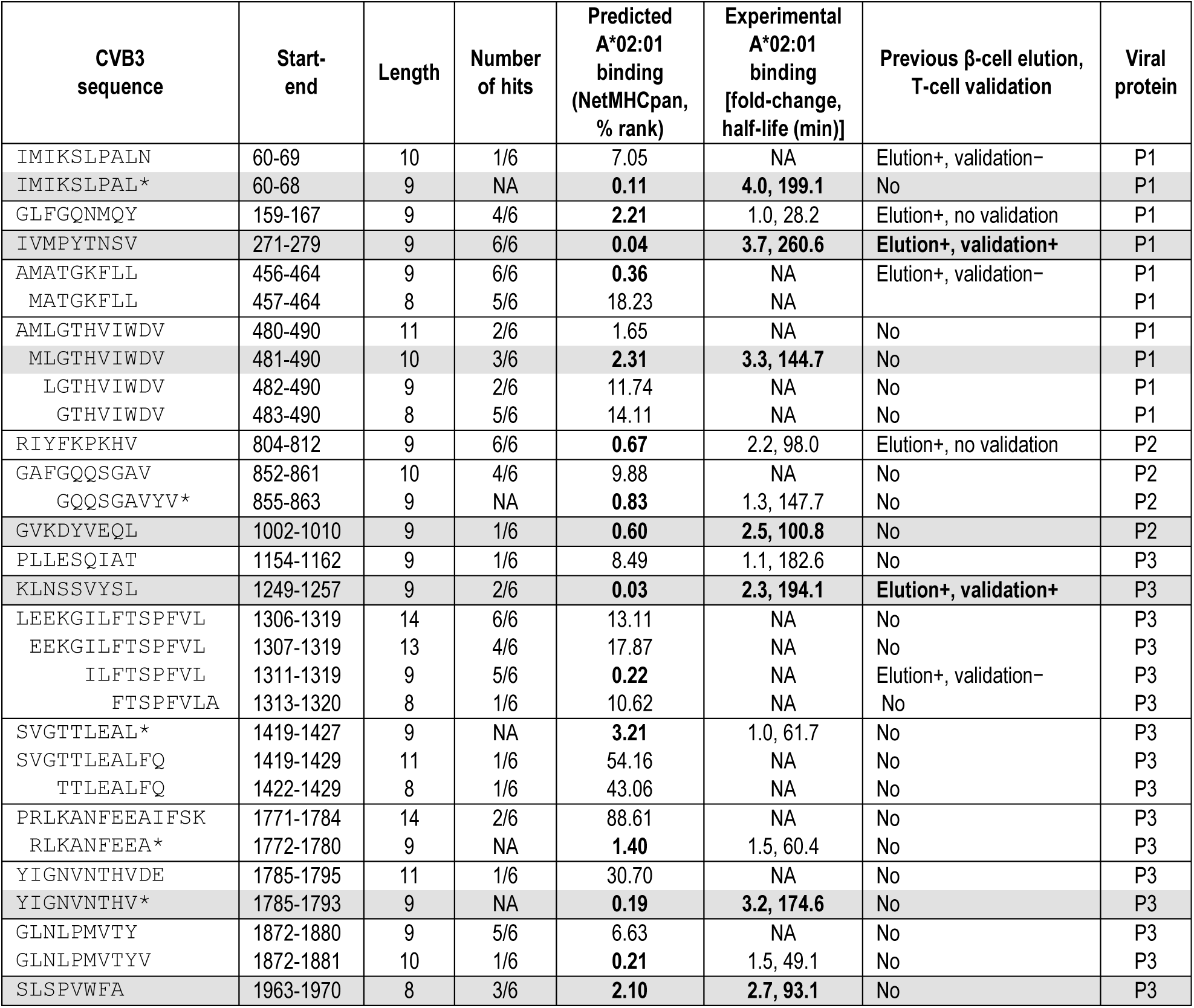
HLA-A*02:01 immunopeptidome eluted from the CaCo2 infection model. For each predicted binder, all overlapping length variants eluted are listed, with asterisks denoting non-eluted length variants with better binding scores. Peptides displaying a predicted NetMHCpan 4.2 binding rank ≤4% (scores marked in bold) were selected for in-vitro HLA-A*02:01 binding assays (using the 9-10mer length variant with the best predicted binding; Fig. S2), except for those that were previously eluted from CVB-infected β cells but did not validate for CD8^+^ T-cell recognition^9^ (second last column). Peptide PLLESQIAT (NetMHCpan rank 8.49%) was included in binding assays as a negative control. HLA-A*02:01 binders were retained when scoring a ≥2.3 fold-change vs. DMSO diluent in in-vitro binding assays (scores marked in bold). Confirmed HLA-A*02:01 peptide binders (highlighted in gray) were further selected for T-cell assays.

Collectively, these results highlight a modest CVB3-mediated cytopathic effect, but no CD8^+^ T-cell-mediated cytotoxicity due to immune escape through downregulation of HLA-I antigen presentation. In contrast, CD4^+^ T-cell responses are amplified through HLA-II upregulation.

### Both direct infection and viral antigen uptake lead to a limited presentation of HLA-bound CVB3 peptides focused on specific viral regions

Given the modulation of HLA-I and HLA-II pathways in CVB3-infected CaCo2 enterocytes, we next aimed to define the HLA-displayed viral peptides using high-resolution mass spectrometry. To this end, we employed two complementary infection models (Fig. 3A). First, CVB3-infected CaCo2 enterocytes to model the primary antigen presentation at the gut entry site. Second, CVB3-infected ECN90 β cells which, upon apoptosis, were allowed to be taken up by DRB1*04:01-transduced THP-1 (DR4/THP-1) monocytic cell line differentiated into macrophages, to model the secondary antigen presentation at the pancreas target site.

**Figure 3.**
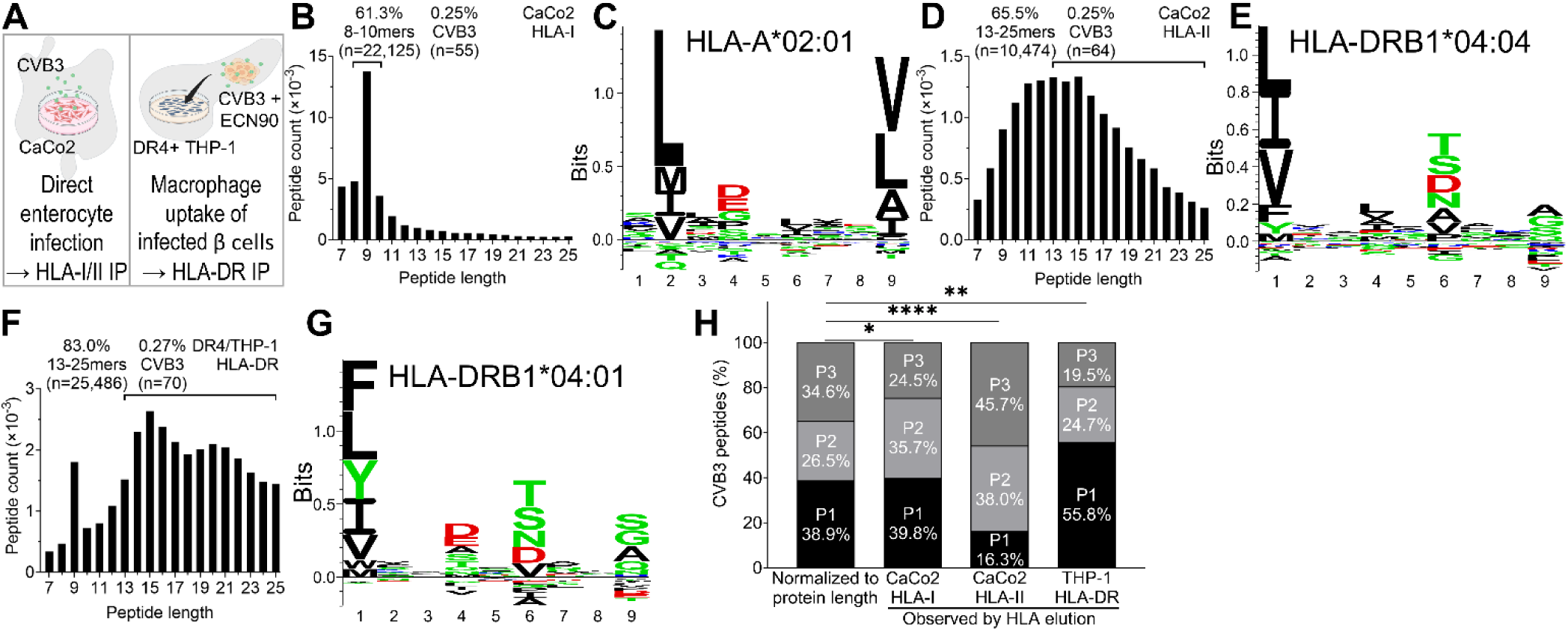
Both direct infection and viral antigen uptake lead to presentation of HLA-bound CVB3 peptides focused on specific viral regions. **A.** Schematic of the in-vitro infection used for immunopeptidomics studies: *left*, direct enterocyte infection (CaCo2 cells, CVB3 300 MOI, 10 h), followed by HLA-I IP (W6/32 Ab; panels B-C) and HLA-II IP (IVA12 Ab; panels D-E); *right*, macrophage uptake of infected β cells [DRB1*04:01-transduced (DR4)/THP-1 macrophages co-cultured with apoptotic CVB3-infected ECN90 β cells, 10 MOI, 72 h], followed by HLA-DR IP (L243 Ab; panels F-G). **B.** Length distribution of HLA-I-bound peptides eluted from CVB3-infected CaCo2 enterocytes. Six biological replicates (227-307×10^6^ cells/each) were acquired, and unique peptides across replicates were counted. Bars represent cumulative peptide counts from all replicates. Number and percentage of expected 8-10mer peptides, and of CVB3-derived peptides withing this length range, are indicated. **C.** Predominant binding motif of eluted unique peptides visualized by MHCMotifDecon, consistent with an HLA-A*02:01-binding motif. The x-axis indicates the residue position within the 9mer core sequence. Each aa is represented by its single-letter code, with its size proportional to its frequency at the indicated position. **D-E.** Length distribution of HLA-II-bound peptides (D) and predominant HLA-DRB1*04:04-like binding motif of eluted unique peptides (E) from CVB3-infected CaCo2 enterocytes. Replicates and data representation are the same as in panel B-C. **F-G.** Length distribution of HLA-DR-bound peptides (F) and predominant HLA-DRB1*04:01-like binding motif of eluted unique peptides (G) from DR4/THP-1 macrophages exposed to CVB3-infected ECN90 β cells. Four biological replicates (14-15×10^6^ cells/each) were acquired, and unique peptides across replicates were counted. Data representation is the same as in panels B-C. **H.** Percent distribution across the structural P1 and non-structural P2 and P3 viral proteins of HLA-eluted unique CVB3 peptides (infected CaCo2 cells; second and third bar; DR4/THP-1, fourth bar) vs. those expected based on the aa length of each protein (first bar). *p=0.048, **p=0.004, ****p>0.0001 by χ^2^ test.

In the primary antigen presentation model of CVB3-infected CaCo2 enterocytes, the HLA-I dataset obtained by immunoprecipitation (IP) with the pan-HLA-I Ab W6/32 yielded the expected 8-10mer peptide length distribution, with a dominant peak at 9 amino acids (aa; Fig. 3B). Of the 22,125 unique 8-10mer sequences eluted, 55 (0.25%) were CVB3-derived. The main peptidome motif was consistent with the canonical HLA-A2 hydrophobic anchor residues at position (P)2 (leucine [L], methionine [M], isoleucine [I], valine [V]) and at the C-terminal P9 (V, L, alanine [A], I) (Fig. 3C). Focusing on HLA-A2-restricted peptides, most (7/13, 54%) eluted CVB3 peptides predicted to bind HLA-A2 (Table 1) and experimentally tested in vitro were confirmed as binders (<u>Fig. S2</u>). Among those, 5 novel HLA-A2-restricted peptides not previously detected in CVB-infected β-cell immunopeptidomes (*9*) were identified and retained for further CD8^+^ T-cell studies: CVB3_60-68_, CVB3_481-490_, CVB3_1002-1010_, CVB3_1785-1793_ and CVB3_1963-1970_. Several HLA-A2 ligands previously eluted from β cells (*9*) were also identified in CaCo2 enterocytes, including 2 previously validated for CD8^+^ T-cell recognition: CVB3_271-279_ and CVB3_1249-1257_ (previously annotated as CVB1_1246-1254_; 100% identity across serotypes).

The HLA-II dataset obtained from CVB3-infected CaCo2 enterocytes by IP with the pan-HLA-II Ab IVA12 displayed the expected length distribution skewed toward peptides ≥13 aa-long (*27*) (Fig. 3D); 64 (0.25%) of the 10,474 unique 13-25mer sequences eluted were CVB3-derived. The main peptidome motif was consistent with an HLA-DRB1*04:04 restriction: P1 harboring hydrophobic (L, I, V, M) or aromatic residue (phenylalanine [F], tyrosine [Y]); polar (threonine [T], serine [S], aspartic acid [D], asparagine [N]) or small residue (A, V) at P6; and heterogeneity at P9 (Fig. 3E).

Similarly, the secondary DR4/THP-1 macrophage antigen presentation model yielded an immunopeptidome (using the anti-HLA-DR Ab L243) enriched in long (≥13 aa) sequences (Fig. 3F). Of the 25,486 unique 13-25mer sequences eluted, 70 (0.27%) were CVB3-derived, with an overall predominant DRB1*04:01 motif (Fig. 3G): similar to DRB1*04:04 at P1 and P6, and with hydrophobic/small (S, glycine [G], A) or polar residue (Q, N) at P9. This high motif similarity between DRB1*04:04 and DRB1*04:01 is consistent with the reported peptide binding promiscuity of these two allotypes (*28*). We therefore also considered predicted DRB1*04:04 binders for downstream HLA-DRB1*04:01 prediction and experimental binding analyses. Most (5/6, 83%) eluted CVB3 peptides predicted to bind HLA-DRB1*04:01 (Table 2) were experimentally confirmed in vitro using T-cell competition binding assays (<u>Fig. S3</u>): CVB3_29-44_, CVB3_256-270_, CVB3_889-902_, CVB3_1431-1445_ and CVB3_1620-1634_.

**Table 2.**
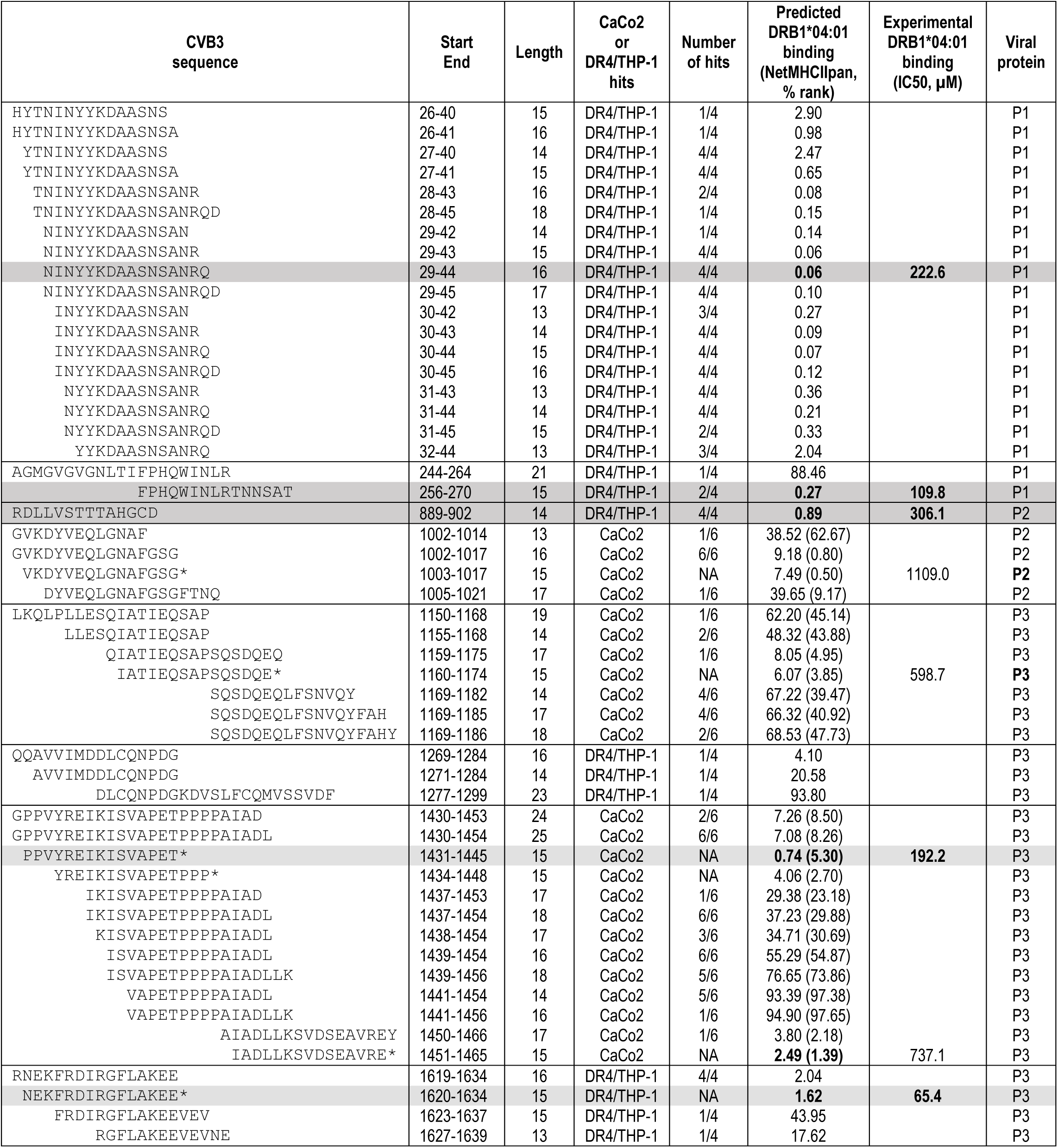
HLA-DRB1*04:01 immunopeptidome eluted from the CaCo2 and DR4/THP-1 infection models. For each predicted binder, all overlapping length variants eluted are listed, with asterisks denoting non-eluted length variants with better binding scores. Peptides displaying a predicted NetMHCIIpan binding rank ≤4% (scores marked in bold; predicted DRB1*04:04 binding in parenthesis for peptides eluted from CaCo2 enterocytes) were selected for in-vitro HLA-DRB1*04:01 binding assays (using the 14-16mer length variant with the best predicted binding). Peptides VKDYVEQLGNAFGSG and IATIEQSAPSQSDQE (NetMHCIIpan rank 7.49% and 6.07%, respectively) were included in binding assays as negative controls. HLA-DRB1*04:01 binders were retained when scoring an IC50<400 µM vs. DMSO diluent in in-vitro binding assays (scores marked in bold). Confirmed HLA-DRB1*04:01 peptide binders (highlighted in gray) were further selected for T-cell assays.

Mapping of the CVB3-derived peptides identified to structural (P1) and non-structural (P2/P3) regions of the viral polyprotein (Fig. 3H) revealed a skewed distribution specific to each in-vitro model. In the primary antigen presentation model of infected CaCo2 enterocytes, the percentage of HLA-eluted peptides mapped significantly more than expected to non-structural regions (based on their relative aa length): P2 for HLA-A2, as we previously observed in ECN90 β cells (*9*); both P2 and P3 for HLA-DRB1*04:04. In contrast, the secondary antigen presentation model of DR4/THP-1 macrophages yielded an enrichment for peptides derived from the structural region (P1) that builds up mature virions, consistent with a dominant mechanism of exogenous uptake of viral material rather than endogenous viral replication. No open-reading frame (ORF)-translated peptides were detected.

Collectively, both CVB3-infected CaCo2 enterocytes and DR4/THP-1 macrophages exposed to infected β cells display a limited repertoire of CVB3-derived peptides via both HLA-I and HLA-II. These peptides are predominantly derived from non-structural viral proteins in infected cells, and from structural proteins upon endocytosis of viral material. This immunopeptidome expands the HLA-A2-restricted one previously identified in CVB-infected β cells (*9*), and provides a novel HLA-DRB1*04:01-restricted immunopeptidome for CD4^+^ T-cell studies.

### CVB3-reactive CD8^+^ T cells display naïve- and exhausted effector/memory-like phenotypes

The panel of HLA-A2-restricted CVB3 candidate epitopes obtained by immunopeptidomics provided a toolkit to track their cognate CD8^+^ T cells. To this end, circulating CD8^+^ T cells from 6 CVB-seropositive HLA-A2^+^ (A*02:01^+^) healthy donors (<u>Table S1</u>) were analyzed using a combinatorial HLA-A2 multimer (MMr) staining strategy (*9, 29–32*) (<u>Fig. S4A-B</u>). The peptide panel included both novel candidates and two epitopes (CVB3_271-279_, CVB3_1249-1257_) identified both herein and previously from CVB-infected β cells, and previously validated for CD8^+^ T-cell recognition (*9*). Control Flu MP_58-66_ and MelanA_26-35_ epitopes were included to provide a benchmark for effector/memory and naïve phenotypes, respectively.

Barring CVB3_60-68_, cognate MMr⁺CD8⁺ T cells were detectable in all donors for the entire set of CVB3 peptides (Fig. 4A; except CVB3_1002-1010_ for donor HD04), thus validating them as target epitopes. However, their frequencies were overall low, with median values ranging from 5 to 70/10^6^ CD8⁺ T cells, i.e. ∼10- to 100-fold lower than what observed for Flu and MelanA epitope-reactive CD8^+^ T cells (median frequency 815 and 445/10^6^ CD8^+^ T cells, respectively). More robust, yet overall weak responses were detected for the novel epitope CVB3_1963-1970_ (median frequency 59/10⁶ CD8⁺ T cells).

**Figure 4.**
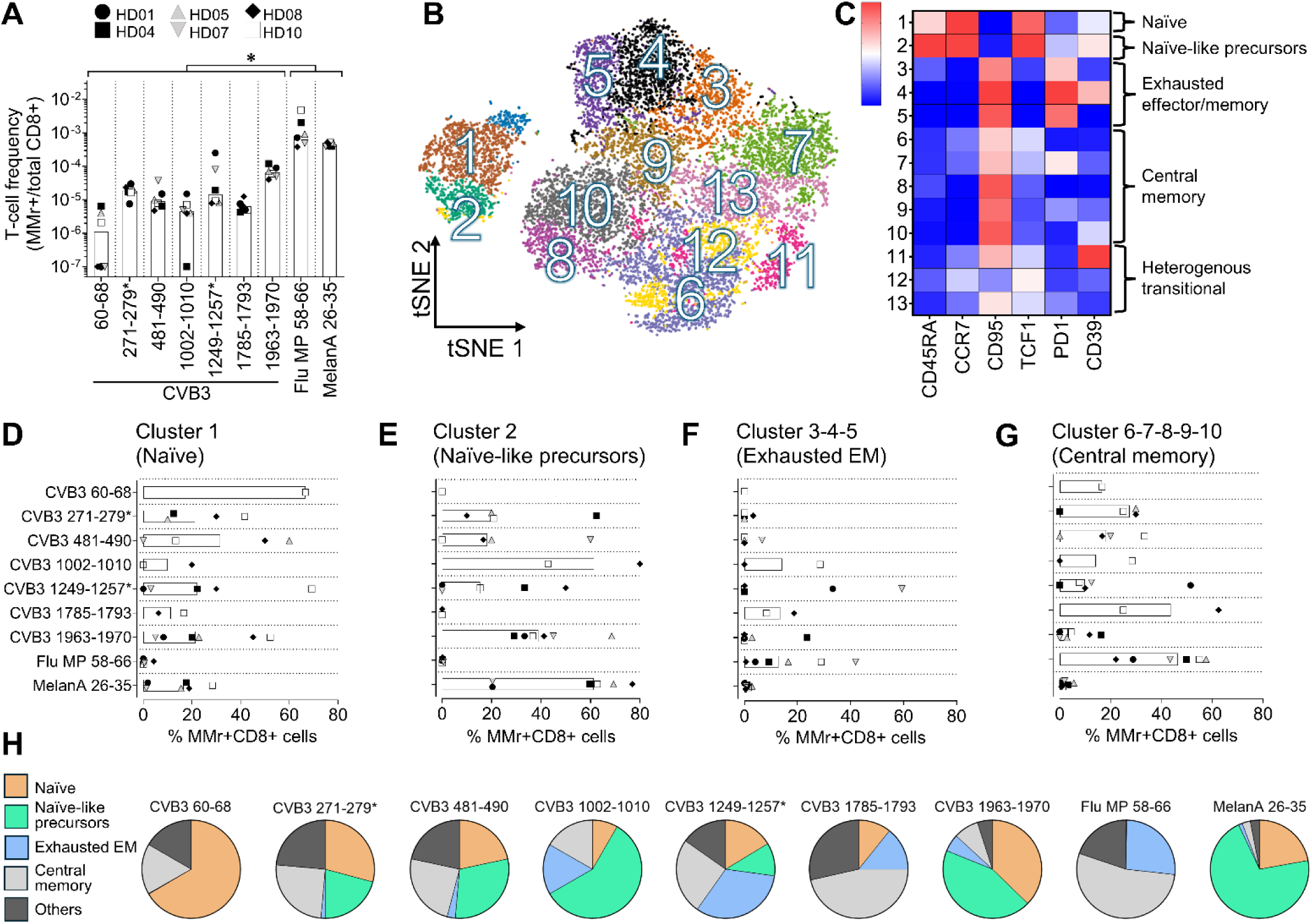
Naïve- and exhausted effector/memory-like phenotypes in CVB3-reactive CD8_+_ T cells. **A.** Blood frequency of MMr^+^CD8^+^ T cells recognizing the indicated CVB3 peptides in CVB-seropositive HLA-A*02:01^+^ healthy donors (n=6). The frequency of Flu MP_58-66_ and MelanA_26-35_ peptide-reactive CD8^+^ T cells included as controls for effector/memory and naïve phenotypes, respectively, is shown for comparison. Data points with <5 MMr^+^ cells counted were excluded. The CVB3_271-279_ and CVB3_1249-1257_ epitopes previously identified in β-cell immunopeptidomes and validated for T-cell recognition are indicated by asterisks. Each symbol represents a donor and bars indicate median values. \**p*=0.031 for the individual comparison of each CVB3 MMr^+^ frequency with Flu or MelanA MMr^+^ frequency by Wilcoxon signed rank test. **B.** tSNE projection of MMr^+^CD8^+^ T cells, clustered based on the expression of the indicated phenotypic markers of T-cell differentiation (CD45RA, CCR7, CD95, TCF1) and activation/exhaustion (PD-1, CD39). Each dot represents a single cell. **C.** Heatmap of normalized expression intensity of each phenotypic marker across the 13 clusters. **D-G.** Percent MMr^+^CD8^+^ T cells across clusters. Each symbol represents a donor and bars indicate median values. **H.** Cumulative relative cluster distribution in MMr^+^CD8^+^ T cells reactive to the indicated peptides.

We next examined the phenotypic composition of these epitope-reactive populations by projecting all MMr⁺CD8⁺ T cells into a t-distributed stochastic neighbor embedding (t-SNE) map generated from multiparametric flow cytometry data (Fig. 4B; representative staining in <u>Fig. S4C</u>). This allowed us to visualize their distribution across 13 clusters defined by differentiation (CD45RA, CCR7, CD95, TCF1) and activation/exhaustion markers (PD-1, CD39) (Fig. 4C). Cluster 1 displayed a naïve phenotype (CD45RA⁺CCR7⁺CD95⁻TCF1⁺). Cluster 2 displayed the same naïve-like homing phenotype, yet co-expressed CD39 and, to a lesser extent, PD-1, indicating precursors with features of chronic activation. Clusters 3-4-5 corresponded to effector/memory T cells (CD45RA^−^CCR7^−^CD95^hi^TCF1^−^) with exhaustion features (PD-1^hi^ with variable CD39 expression). In comparison, clusters 6 to 10 re-expressed some CCR7 and were CD95^int^TCF1^lo^, corresponding to central memory CD8^+^ T cells. Clusters 11-13 had more heterogenous, transitional phenotypes.

The distribution of epitope reactivities within these clusters revealed distinct patterns (Fig. 4D-H). Barring CVB3_60-68_ due to its low cognate T-cell frequencies, most epitope reactivities (CVB3_271-279_, CVB3_481-490_, CVB3_1002-1010_, CVB3_1963-1970_) were dominated by naïve and naïve-like precursors, while CVB3_1249-1257_ featured higher proportions of exhausted effector/memory CD8^+^ T cells. The representation of central memory clusters was overall low across all CVB3 reactivities, barring their unique dominance for the novel CVB3_1785-1793_ epitope, yet also recognized at low frequencies (Fig. 4A). The control Flu MP_58-66_- and MelanA_26-35_-reactive CD8^+^ T cells displayed the expected predominant central memory and naïve-like phenotypes, respectively.

Collectively, while some CVB3-reactive CD8^+^ T cells progress from a naïve to an activated naïve-like phenotype, this progression is limited by the development of exhaustion features, preventing their differentiation into central memory subsets.

### CVB3-reactive CD4^+^ T cells exhibit robust memory and Tfh phenotypes

The limited CVB3 antiviral responses measured in the CD8^+^ T-cell compartment beg the question of whether these responses may rely predominantly on CD4^+^ T cells. To address this possibility, CD4⁺ T-cell responses were evaluated in CVB-seropositive HLA-DRB1*04:01^+^ healthy donors (<u>Table S1</u>), using the HLA-DRB1*04:01-restricted peptides identified by immunopeptidomics and the Flu HA_306-318_ epitope as a benchmark for effector/memory responses. Following a 24-h stimulation of peripheral blood mononuclear cells (PBMCs) with each individual peptide, CD4^+^ T cells were analyzed by flow cytometry for the expression of activation-induced marker (AIM) signatures (*33*), comprising CD40L, 4-1BB, CD25, CD69, HLA-DR, OX40, and PD-L1. For each donor/peptide combination, after gating on activated CD40L^+^CD4^+^ T cells (<u>Fig. S5A</u>) all 64 possible AIM combinations were assessed (i.e. CD40L alone or in combination with 1, 2, 3, 4, 5 or 6 other AIMs; <u>Fig.</u> <u>S5B</u>). To achieve maximal specificity and sensitivity, the combinations retained for each donor were those that *a)* did not yield any AIM^+^CD4^+^ T cell following the DMSO negative control stimulation; and *b)* did yield AIM^+^CD4^+^ T cells following each peptide stimulation analyzed individually (<u>Fig. S5C-D</u>).

In all donors, CVB3-reactive AIM⁺CD4⁺ T cells were readily detectable for all peptides tested, generally clustering together within a 10^−3^ frequency range (i.e., ∼1,000 peptide-reactive T cells per million CD4⁺ T cells), comparable to the frequency measured for the recall control Flu HA_306-318_ epitope (Fig. 5A). While donors HD09 and HD10 displayed lower frequencies, the magnitude of CVB3-reactive responses also paralleled that for Flu HA_306-318_ in donor HD10. Overall, this validates the CVB3 peptides selected by immunopeptidomics as epitopes robustly targeted by CD4^+^ T cells.

**Figure 5.**
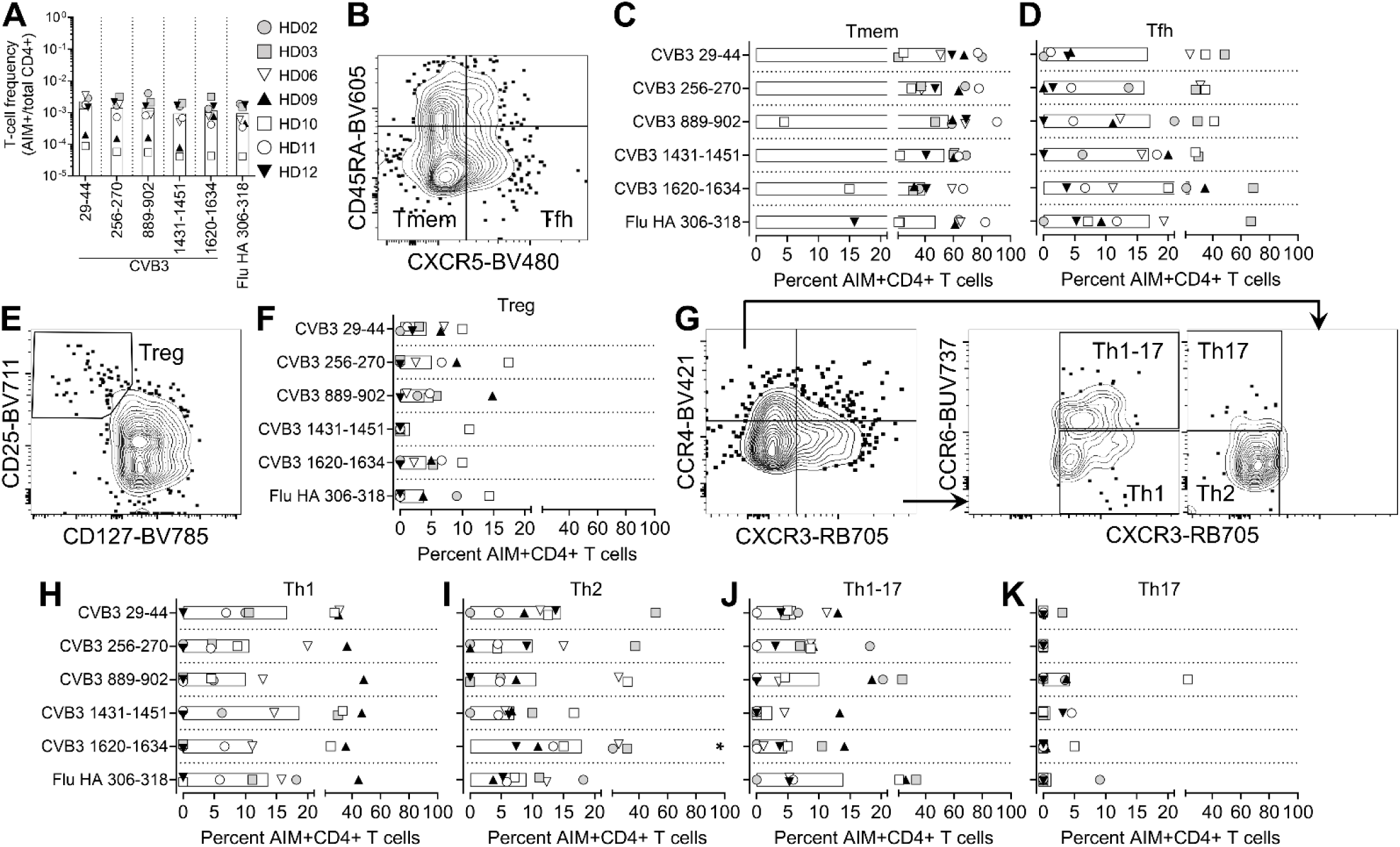
CVB3-reactive CD4_+_ T cells exhibit robust memory and Tfh phenotypes. **A.** Blood frequency of AIM^+^CD4^+^ T cells recognizing the indicated CVB3 peptides after a 24-h peptide stimulation in CVB-seropositive HLA-DRB1*04:01^+^ healthy donors (n=7). The frequency of Flu HA_306-318_-reactive CD4^+^ T cells included as effector/memory control is shown for comparison. **B.** Gating strategy to identify effector/memory (Tmem, CD45RA^−^CXCR5^−^) and follicular helper T cells (Tfh; CD45RA^−^CXCR5^+^) subsets within AIM^+^CD4^+^ T cells. **C-D.** Percentages of Tmem (C) and Tfh (D) subsets within AIM^+^CD4^+^ T cells reactive to each peptide. **E.** Gating strategy to identify regulatory T cells (Treg, CD25^high^CD127^low^) within AIM^+^CD4^+^ T cells. **F.** Treg percentages within AIM^+^CD4^+^ T cells. **G.** Gating strategy to identify T helper (Th) polarization based on CCR4, CXCR3, and CCR6 expression within AIM^+^CD4^+^ T cells: Th1 (CXCR3^+^CCR4^−^CCR6^−^), Th2 (CXCR3^−^CCR4^+^CCR6^−^), Th17 (CXCR3^−^CCR4^+^CCR6^+^) and Th1/Th17 (CXCR3^+^CCR4^−^CCR6^+^). **H–K.** Percentages of Th1 (H), Th2 (I), Th1/Th17 (J) and Th17 (K) subsets within AIM^+^CD4^+^ T cells. Each symbol represents a donor and bars indicate mean values. *p=0.016 for pairwise comparison with Flu HA-reactive AIM^+^CD4^+^ T cells by Wilcoxon signed rank test.

To define functional phenotypes, AIM⁺CD4⁺ T cells were first analyzed for their CD45RA and CXCR5 expression to define conventional memory T cells (Tmem; CD45RA⁻CXCR5⁻) and follicular helper T cells (Tfh; CD45RA⁻CXCR5⁺) (Fig. 5B). Overall, more than 40% of CVB3-reactive CD4⁺ T cells displayed a Tmem phenotype, again comparable to that of control Flu HA-reactive CD4^+^ T cells (Fig. 5C). Tfh cells accounted for an average of more than 16% of epitope-reactive CD4⁺ T cells for both CVB3 and Flu HA (Fig. 5D). In contrast, regulatory T cells (Tregs; CD25^hi^CD127^lo/neg^; Fig. 5E) were less frequent (<5%) in most instances (Fig. 5F), and their fractions were inversely correlated with T-cell frequencies (Spearman r=−0.412, p=0.007; <u>Fig. S5E</u>). These findings suggest that CVB3-reactive CD4⁺ T cells mount a robust memory and provide effective Tfh support, enabling the generation and maintenance of antiviral antibody responses.

We expanded our analysis to determine the T helper (Th) polarization of AIM⁺CD4⁺ T cells. Th subsets were defined using CCR4, CXCR3, and CCR6 expression (Fig. 5G): Th1 (CXCR3⁺CCR4⁻CCR6⁻), Th2 (CXCR3⁻CCR4⁺CCR6⁻), Th17 (CXCR3⁻CCR4⁺CCR6⁺), and Th1/Th17 (CXCR3⁺CCR4⁻CCR6⁺). All CVB3 peptides elicited detectable Th1, Th2 and Th1/Th17 responses in most but not all donors (Fig. 5H-I<u>-J</u>), with no statistically significant donor-paired differences compared to Flu HA. The only exception was CVB_1620-1634_, which showed a higher Th2 polarization compared to Flu HA. Th17 frequencies were low or undetectable across all epitopes (Fig. 5K).

Collectively, these results show that CVB3-reactive CD4⁺ T cells mount robust, polyfunctional memory responses, comparable to those elicited by Flu HA. This capacity contrasts with the limited effector/memory differentiation of CVB3-reactive CD8⁺ T cells.

## Discussion

Understanding how immunity against CVB3 is shaped at the intestinal entry site could provide crucial insights into the mechanisms driving the onset of islet autoimmunity in T1D. This standing question has direct clinical relevance, as multiple studies have detected enteroviral persistence in the gut of T1D patients, implicating early mucosal immune responses as potential triggers of downstream β-cell damage (*4, 8, 15*). We addressed this question by focusing on antigen presentation, the main gatekeeper of T-cell-mediated immunity. While our previous work (*9*) focused on the dynamics of T-cell-mediated cytotoxicity following viral antigen presentation by infected β cells, in the present work we considered both the earliest intestinal location of viral entry and the later events following death of infected β cells and uptake by macrophages. To this end, we employed two complementary infection models that recapitulate these routes of antigen presentation: an enterocyte model of direct infection, and a macrophage model of indirect antigen uptake. These models combined the analysis of the host proteome, HLA-bound viral peptidome and the ensuing T-cell responses to provide a comprehensive view of antiviral immune recognition.

At the proteomic level, we observed remodeling of multiple pathways that collectively impair antigen processing and presentation, alongside broad suppression of innate immune sensing. These changes are in agreement with the known activity of viral proteases (*34*), which cleave key adaptor proteins in the RIG-I-like and Toll-like receptor signaling pathways (*35*), disrupt ER-to-Golgi trafficking, and hinder HLA-I peptide loading (*36–38*). CVB3 also halted cell proliferation, which may redirect host biosynthetic resources toward viral replication (*39*), lowering protein turnover and, in turn, immune recognition (*40, 41*). Alterations in autophagy-related proteins suggested increased early autophagic events but inhibition of late maturation and lysosomal fusion steps, a pattern reported for other Enteroviruses that exploit autophagy to generate replication membranes while restricting the degradative flux (*42*). As in β cells (*9*), live-cell imaging further revealed filopodia-mediated viral transmission from infected enterocytes. Non-lytic CVB shedding has been previously reported, either via membrane protrusions or autophagosome-derived vesicles, both of which can shield virions from immune recognition and Ab-mediated neutralization (*43, 44*). These findings point to multiple, converging CVB3 immune escape strategies.

Guided by these proteomic signatures, surface protein expression and functional assays confirmed a reduction of surface HLA-I expression. This resulted into downstream inhibition of antiviral CD8^+^ T-cell recognition and impaired killing of infected CaCo2 enterocytes, which were instead killed when the cognate CVB3 peptide was exogenously added. This suggests that CVB3 specifically limits exposure of its own peptides on top of HLA-I downregulation. Limited T-cell cytotoxicity was also previously observed in β cells, which were rather killed by the viral cytopathic effects (*9*), suggesting a conserved immune evasion mechanism across different target tissues. Unexpectedly, infected enterocytes further revealed an upregulation of HLA-II, which resulted in an opposite outcome of enhancing CD4^+^ T-cell recognition. Thus, the net effect of direct antigen presentation in infected enterocytes is a viral escape from CD8⁺ T-cell cytotoxic surveillance and preservation of CD4⁺ T-cell responses.

Looking at the CVB3 antigen presentation landscape, the HLA-A2- and HLA-DRB1*04:01-restricted immunopeptidome displayed by enterocytes was narrow and predominantly composed of non-structural proteins, as reported for the HLA-I immunopeptidome of infected β cells (*9*). This may be expected in the context of productive infection and active intracellular viral replication, because non-structural proteins do not leave the cell with the release of mature virions but are eventually catabolized intracellularly, thus providing more accessible substrates for antigen presentation. In contrast, the HLA-DRB1*04:01-restricted immunopeptidome eluted from the non-replicative macrophage model was focused on structural proteins, consistent with the uptake and processing of intact virions and viral debris. From these datasets, we validated 5 HLA-I-eluted and 5 HLA-II-eluted CVB3 peptides as HLA-A2- and HLA-DRB1*04:01-binding epitopes targeted by CD8^+^ and CD4^+^ T cells, respectively. This panel further included 2 epitopes (CVB3_271-279_ and CVB3_1249-1257_) eluted from both infected enterocytes and β cells and previously validated as the immunodominant targets of CD8^+^ T cells (*9*). These reference epitopes were recognized by CD8^+^ T cells at low frequencies, similar to those observed for the novel ones identified.

This more comprehensive epitope panel afforded greater granularity to analyze anti-CVB3 cellular immunity than previously obtained from the β-cell peptidome, notably by tracking both CD8^+^ and CD4^+^ T cells. Expanding our previous observation (*9*) of high circulating naïve CD8⁺ T-cell fractions, we here confirm that CVB3-reactive CD8^+^ T-cell responses display low frequencies, reflecting early differentiation states comprising both naïve- and effector/memory-like phenotypes with exhaustion features. These phenotypes included a pool of naïve-like precursors that retain lymphoid homing markers but have acquired chronic activation/exhaustion markers (PD-1, CD39), likely as a result of repeated or persistent exposure to CVB antigens. On one hand, this predominance of circulating early-differentiated intermediates may reflect the preferential retention of effector/memory fractions within lymphoid organs (*9*). On the other, suboptimal CD8⁺ T-cell priming and early exhaustion may result from the CVB3 immune escape mechanism of downregulating HLA-I antigen presentation. This may ultimately compromise viral clearance by CD8⁺ T cells and favor the reported CVB persistence (*3, 7*).

In contrast, DRB1*04:01-restricted CVB3 epitopes were consistently immunogenic: they activated a considerable fraction of memory CD4^+^ T cells in most donors, including a sizable Tfh subset. This activation was broadly polarized towards Th1, Th1/17 and Th2, but not Th17, responses. All these features were similar to those observed for the prototypic viral recall epitope Flu HA_306-318_, suggesting that protective CD4^+^ T-cell responses are efficiently mounted and maintained after CVB infection. Beyond supporting cytotoxic responses, CD4^+^ T cells orchestrate broader antiviral defenses (*45*), inducing humoral responses and cross-talking with innate-like lymphocytes. While the Th1 arm supports IFN-γ-mediated APC maturation and CD8⁺ T-cell cytotoxicity, Tfh and Th2 responses promote B-cell germinal center and Ab maturation (*45*). Hybrid Th1/17 differentiation may instead be a double-edged sword (*46*): while it can aid mucosal viral clearance via IL-17-mediated innate cell recruitment, increased IL-17 with Th1-skewed inflammation has also been linked to immunopathology, including in CVB3 infection (e.g., myocarditis) (*47, 48*). Tight regulation of Th1/17 skewing may thus be needed to secure viral immunity while minimizing collateral inflammation. Altogether, these results indicate that antiviral CVB control may predominantly rely on the CD4⁺ T-cell arm, compensating the limited cytotoxic CD8^+^ T-cell responses. Accordingly, a curated compilation from the Immune Epitope Data Base (IEDB) (*49*) shows that, in Enteroviruses, the few HLA-I-restricted epitopes identified map mainly to non-structural proteins and elicit low CD8^+^ T-cell response rates, whereas HLA-II-restricted epitopes are more numerous, map largely to structural proteins, and elicit higher CD4^+^ T-cell response rates. This is consistent with previous reports of CD4^+^ T-cell responses focused on structural proteins also for CVB4 (*50*).

Triggering a CD8-deficient T-cell immunity may thus be a general strategy mounted by Enteroviruses to avoid cytotoxic clearance of infected cells. While this limited CD8^+^ T-cell immune pressure may favor low-grade viral persistence, the efficient CD4^+^ T-cell responses provide therapeutic opportunities to harness them toward protective antiviral immunity. On one side, this may limit CD8^+^ T-cell exhaustion, on the other it may amplify a productive cross-talk with B cells and innate-like lymphocytes. CD4⁺ T cells can coordinate with mucosal-associated invariant T (MAIT) cells (*51, 52*) and invariant natural killer T (iNKT) cells (*53*) to stimulate germinal centers. In parallel, innate lymphoid cells (ILCs, particularly ILC3 and ILC2) shape the cytokine milieu that instructs Th polarization and mucosal B-cell class switching, reinforcing a feed-forward antiviral loop (*54*). Against this background, a mucosal CVB vaccine may boost more robust CD4^+^ T-cell responses, notably Tfh, promoting IgA secretion at the gut entry site while recruiting innate-like helpers to accelerate and stabilize protective immunity and compensate the escape from CD8^+^ T cells. Evidence from other viruses shows that mucosal vaccination elicits tissue-restricted immunity that systemic vaccines often miss. For instance, intranasal live attenuated influenza vaccines induce robust nasal IgA and CD4^+^ T-cell responses (*55*). Expanding CVB antigen coverage to non-structural proteins may also overcome the HLA-I “visibility” bottleneck at the gut entry site. As CVB vaccination strategies move into clinical trials (*13*), our findings may inform the design of next-generation candidates, e.g. live attenuated oral CVB vaccines boosting mucosal immunity, as successfully done with the nOPV2 vaccine against the CVB-related poliovirus (*56, 57*). These vaccination strategies may be complemented by antiviral therapies (*10*), e.g. the capsid-binding compound pleconaril and enteroviral 2C protein inhibitor fluoxetine (*58*) that may reduce CVB-mediated suppression of IFN and HLA-I pathways. Our validated HLA-I and HLA-II epitope panels also provide immune monitoring tools to track the impact of these interventions on CD8⁺ and CD4⁺ T cells.

Limitations of this study include the use of reductionist in-vitro models that may not fully capture the in-vivo setting. However, our findings are in line with mouse studies of in-vivo CVB3 infection documenting poor CD8^+^ and robust CD4^+^ T-cell responses (*59*). Second, we do not provide formal proof that THP-1 macrophages exposed to infected β cells are not in turn infected by CVB3. Mitigating this potential, macrophages are reportedly resistant to CVB infection (due to low expression of the CAR CVB entry receptor and efficient type I IFN responses leading to abortive replication) (*60*). Moreover, the observed peptide display was focused on structural rather than non-structural CVB3 regions, consistent with a predominant antigen uptake mechanism. Third, our T-cell analyses were limited to healthy donors, preventing causal inference for T1D, i.e. whether these responses are impaired in individuals later developing disease. Addressing this gap will require longitudinal tracking of CVB-reactive CD4⁺/CD8⁺ T-cell trajectories during and after natural infection.

In conclusion, CVB3-driven immune escape mechanisms at the intestinal entry site impair antiviral CD8⁺ T-cell immunity by stalling precursors on naïve and exhausted effector/memory differentiation states. In contrast, CVB3-reactive CD4⁺ T cells differentiate into polyfunctional memory helper and Tfh subsets. These results lend rationale to mucosal vaccination strategies to boost anti-CVB immunity more efficiently and provide novel tracking tools to follow T-cell modifications following natural infection or vaccination.

## Materials and Methods

### Experimental design

The objective of this study was to understand the dynamics driving antiviral CVB immunity at the intestinal entry site. Initial proteome analysis of CVB3-infected enterocytes revealed several immune escape mechanisms, including dysregulation of antigen processing and presentation pathways. Following on this observation, we explored the surface expression of HLA-I and HLA-II upon CVB3 infection and downstream modulation of CD4^+^ and CD8^+^ T-cell responses. These functional experiments highlighted escape mechanisms from CD8^+^ T cells. In contrast, in-vitro CD4^+^ T-cell activation was enhanced by HLA-II upregulation. These findings set the stage for our hypothesis that CVB3 infection in the primary replication site of enterocytes drives limited peripheral CD8^+^ T-cell immunity but more robust CD4^+^ T-cell responses. To track these CVB3-reactive CD4^+^ and CD8^+^ T-cell responses, we identified HLA-I- and HLA-II-bound peptides from two in-vitro models of CVB3 antigen presentation: replicative enterocyte infection and macrophage uptake of viral debris. Despite limited viral antigen presentation, we established an HLA-A2- and HLA-DRB1*04:01-restricted candidate epitope panel comprising non-structural CVB3 peptides, over-represented in our replicative enterocyte model; and structural CVB3 peptides, enriched in our macrophage uptake model. This panel allowed us to track the frequency and phenotype of circulating CVB3-reactive CD8^+^ T cells, using ex-vivo HLA-A2 MMr assays; and CD4^+^ T cells, using AIM assays upon in-vitro peptide stimulation. This analysis validated our hypothesis: CVB3-reactive CD8^+^ T cells were developmentally stalled in naïve and exhausted effector/memory states, whereas CD4^+^ T cells comprised a broad range of Tfh, Th1, Th2 and Th1/17 phenotypes. All in-vitro experiments were performed at least in triplicates on at least two separate occasions. Blood samples were processed in batch, and no outliers were excluded; only undersampled data points were excluded if necessary.

### Preparation and titration of CVB3 stocks

Viral stocks were prepared as previously described (*9*). Briefly, CVB3 (Nancy strain) was purified from infected HeLa cells, titrated on plaque assays and sequenced. CVB3-GFP was generated by transfection of HeLa cells with a construct generated by cloning the eGFP gene into a previously engineered SfiI site immediately located downstream of the start codon of the CVB3 polyprotein within the pMKS1 plasmid backbone (*61*).

### In vitro infection of CaCo2 enterocytes

The first model of CVB3 antigen presentation employed the enterocyte cell line CaCo2 (RRID: CVCL_0025), cultured in Advanced DMEM/F-12 (ThermoFisher #12634028) supplemented with 2% fatty-acid-free bovine serum albumin (BSA, fraction V; Roche #10775835001), 1% GlutaMax (ThermoFisher, #35050061), 0.02 nM sodium selenite (Sigma, #S9133), 10 mM nicotinamide (Millipore, #481907), 50 µM 2-mercaptoethanol (Sigma, #M6250) and 1% penicillin/streptomycin (ThermoFisher, #15140122). Cells with early passage numbers (25 to 29) were seeded in 150-cm^2^ tissue culture flasks (TPP, #90151) at 3.2×10^6^ cells/flask and cultured for 3 days or until 60-70% confluent. After gentle rinsing with Dulbecco’s phosphate-buffered saline (DPBS), CaCo2 cells were infected with CVB3 (300 MOI) for 1 h in complete medium without BSA, followed by transfer into fresh pre-warmed BSA-free medium. After continuing culture for 10 h, cells were harvested by gentle scraping in ice-cold dissociation buffer (DPBS, 5 mM EDTA). Cell pellets were washed twice in DPBS and split for flow cytometry analysis or freezing at −80°C as dry pellets for downstream proteomics and immunopeptidomics. For proteomics, 9 replicates (5×10^6^ cells/each) were generated: 4 CVB-infected and 5 mock-infected. For immunopeptidomics, 6 replicates (3×10^8^ CVB3-infected CaCo2 cells/each) were processed. Flow cytometry analysis of CaCo2 cells was performed on a BD LSR Fortessa II cell analyser. Surface staining was performed using the monoclonal antibody (mAb) W6/32 to HLA-A/B/C/E (RRID: AB_2917755) and Live/Dead Violet viability marker (ThermoFisher, #L34955). Intracellular staining was performed with primary mAbs to VP1 (clone 3A6) (*62*) and dsRNA (RRID:AB_2651015), secondary Abs mouse anti-rat IgG (IgG; RRID:AB_465229) and AlexaFluor (AF)594-conjugated goat anti-mouse IgG (RRID:AB_2762825), and the eBioscience FoxP3 Fix/Perm kit (ThermoFisher, #5523).

RT-qPCR for *HLA-A*, *HLA-B*, *HLA-C* and *HLA-DRB1*04* genes was performed on CVB3-and mock-infected CaCo2 cells, as previously described (*63*). GAPDH was used as the housekeeping gene, and relative quantification was calculated using the 2^−ΔΔCt^ method.

To assess surface CD107a expression, CaCo2 enterocytes were seeded at 30,000 cells/well in 96-well flat-bottom plates and cultured for 12 h before infection with CVB3-GFP (100 MOI, 1 h) in serum-free DMEM. ECN90 β cells were treated similarly but seeded in plates coated with 0.25% fibronectin from human plasma and 1% extracellular matrix from Engelbreth-Holm-Swarm murine sarcoma, and cultured in DMEM/F12 Advanced medium supplemented with 2% BSA, 50 μM 2-mercaptoethanol, 10 mM nicotinamide, sodium selenite (1.7 ng/ml; Sigma), penicillin/streptomycin and 1% GlutaMAX before infection with CVB3-GFP (50 MOI, 1 h) in serum-free medium and culture for an additional 16 h. After gentle washing in DPBS, the culture was continued (20 h for CaCo2, 16 h for ECN90), followed by trypsinization, washing and processing for flow cytometry. Infection was verified as GFP positivity on viable cells stained with Live/Dead Far Red (ThermoFisher, #L10120) and surface staining with BV786-conjugated mAb to CD107a (RRID:AB_2738458). The median fluorescence intensity (MFI) of CD107a between infected vs. non-infected cells was compared to evaluate the increase in surface CD107a expression.

### In vitro uptake of infected ECN90 β cells by DR4/THP-1 macrophages

The second model of CVB3 antigen presentation employed the macrophage-differentiated DR4/THP-1 cell line exposed to apoptotic, CVB3-infected ECN90 β cells. THP-1 cells (RRID: CVCL_0006) were lentivirally transduced to stably express HLA-DRB1*04:01 and cultured in RPMI-1640 (ThermoFisher, #1870036) supplemented with 20% heat-inactivated fetal bovine serum (FBS; PAN Biotech, #P30-3306) and 1% penicillin/streptomycin. For differentiation, cells were seeded at 0.5×10⁶ cells/mL in 150-cm^2^ flasks and treated with 100 ng/mL phorbol 12-myristate 13-acetate (PMA; Sigma) for 48 h (*64*). Following medium removal, cells were gently rinsed once with pre-warmed DPBS, and fresh PMA/FBS-free medium was added for an additional 24 h rest period to allow recovery and stabilization of the adherent macrophage-like phenotype (*65*), followed by a 24-h co-culture with apoptotic CVB3-infected ECN90 β cells (RRID: CVCL_VJ07) at a 1:1 THP-1/ECN90 ratio. ECN90 β cells were driven to apoptosis by CVB3 infection (10 MOI, 72 h) (*9*). Pulsed DR4/THP-1 cells were harvested by gentle scraping in ice-cold dissociation buffer, washed twice in DPBS and frozen at −80°C as dry pellets for downstream immunopeptidomics. A total of 4 replicates (15×10^6^ cells/each) were processed.

### Proteomics

Frozen cell pellets for proteomics and immunopeptidomics experiments were thawed in lysis buffer (0.5 ml/10⁸ cells) for 30 min on ice. The buffer consisted of 0.5% IGEPAL CA-630 (Sigma, #I8894), 150 mM NaCl, 50 mM Tris-HCl (pH 8.0), and protease inhibitor cocktail (Roche, #11836170001). Lysates were centrifuged at 500g for 10 min at 4°C to remove nuclei, followed by 20,000g for 45 min at 4°C to pellet residual cell debris.

Lysates were quantified using the Pierce BCA Protein Assay (ThermoFisher, #23225) and adjusted to 1 µg/µl in 5% SDS. For each sample, 20 µg protein was reduced with 10 mM DTT for 15 min, alkylated with 55 mM iodoacetamide for 15 min, then digested using the S-Trap micro spin column protocol (ProtiFi). Briefly, proteins were acidified to 2.5% (v/v) phosphoric acid, applied to the S-Trap column, washed with 50 mM TEAB in 90% (v/v) methanol, and digested on-column with 2 µg sequencing-grade trypsin (Promega, #V5111) at 47°C for 1 h. After washing, peptides were eluted with 50% (v/v) acetonitrile (ACN), 0.1% (v/v) formic acid, vacuum-dried, and resuspended in 20 µl 0.1% (v/v) trifluoroacetic acid (TFA), 1% (v/v) ACN.

Peptides were separated on an Ultimate 3000 RSLCnano system (ThermoFisher) using a PepMap C18 EASY-Spray column (75 µm × 50 cm, 5 µm) in 0.1% formic acid and coupled to a Q Exactive HF-X mass spectrometer (ThermoFisher). A 2-35% acetonitrile gradient in water containing 1% DMSO, 0.1% formic acid was run at 250 nl/min. Ionization was via EasySpray at 2,000 V with a transfer tube temperature of 250°C. Data were acquired in proteomics DDA mode: a full MS1 scan (m/z 320-1600; resolution 60,000; AGC 3×10⁶, maximum injection time 45 ms) followed by 12 MS/MS scans (resolution 30,000; AGC 5×10⁵, maximum injection time 54 ms) using a 1.3 m/z isolation width, +2 to +4 charge states, normalized HCD energy 28%, and 30 s dynamic exclusion. All data were recorded in profile mode.

Raw MS data were processed with FragPipe v22.0 (LFQ-MBR workflow) using the same database as for immunopeptidomics (see below). Searches were performed with trypsin specificity (up to two missed cleavages), a precursor mass tolerance of ±20 ppm, and a fragment mass tolerance of ±20 ppm. Carbamidomethylation of cysteine (+57.02146 Da) was set as a fixed modification, while methionine oxidation and N-terminal acetylation were set as variable modifications. Peptide-spectrum matches were filtered to 1% PSM-level FDR with Percolator, and protein inference was performed with ProteinProphet at 1% protein-level false discovery rate (FDR). Label-free quantification was conducted with IonQuant (*66*) (MaxLFQ enabled, match-between-runs active, m/z tolerance 10 ppm, RT tolerance 0.4 min) with normalization across runs. MSBooster (*67*) was also applied.

Label-free quantification (MaxLFQ intensity) data were analyzed in FragPipe-Analyst (*68*). Proteins with ≥70% non-missing values were retained, and missing values were imputed using the Perseus-type method. Differential expression was assessed with Limma, applying a Benjamini-Hochberg-adjusted p≤0.05. Functional enrichment (GO Biological Process, KEGG, Reactome) guided manual selection of proteins related to enterocyte differentiation, antigen processing and presentation, IFN signaling and autophagy.

### Functional T-cell priming assays

To assess the effect of HLA-II upregulation on antigen presentation, CaCo2 cells were infected with CVB3 (10 MOI, 18 h), followed by pulsing with the HLA-DR4-restricted Flu HA_306-318_ peptide (PRYVKQNTLKLAT; 10 µM) or DMSO diluent for 1 h. CaCo2 cells were then co-cultured at a 1:1 ratio with a Flu HA-reactive CD4⁺ T-cell clone for 3 h, after which brefeldin A (BFA, 5 µg/mL; BioLegend #420601) was added for an additional 1 h. Cells were stained with Live/Dead Far Red, followed by intracellular staining with FITC-conjugated TNF-α mAb (RRID:AB_315258). TNF-α expression in CD4⁺ T cells was measured by MFI as a readout of T-cell activation.

To assess the effect of HLA-I downregulation on antigen presentation by CVB3-infected CaCo2 cells, CD8⁺ T-cell-mediated killing was assessed in real time using the Incucyte S3 Live-Cell Analysis System (Sartorius). A stable CaCo2 cell line expressing the NucLight Red nuclear label (mKate2; Sartorius, #4717) was generated by lentiviral transduction following manufacturer’s instructions. Cells (20,000/well, 96-well flat-bottom plates; TPP #92096) were infected with CVB3-GFP at 10 MOI. After 1 h, unbound virions were removed by washing with pre-warmed medium, and fresh medium was added. CVB3_1246–1254_ TCR-transduced primary CD8⁺ T cells (*9*) were then added at a 1:1 effector-to-target ratio. Control conditions included: *i)* infected CaCo2 cells without CD8⁺ T cells, *ii)* infected CaCo2 cells with irrelevant CD8⁺ T cells transduced with the PPI_15-24_ TCR 1E6 (*9*), *iii)* infected CaCo2 cells pulsed with CVB3_1246–1254_ peptide (1 µM) and co-cultured with CVB3-reactive CD8⁺ T cells, and *iv)* mock-infected CaCo2 cells co-cultured with CVB3-reactive CD8⁺ T cells. Plates were scanned every 2 h for 68 h with phase contrast, green (GFP; acquisition time 300 ms), and red (mKate2 Red; acquisition time 400 ms) channels at 10× magnification. Green and red object masks were generated using surface fit segmentation with a threshold of 2 calibration units. Fold changes in CaCo2 cell numbers were quantified as the red object count per image normalized to the value at 0 h. Percentages of infected cells were determined by calculating the ratio of green and red area overlap to the total red area.

### Immunopeptidomics

Before IP of pHLA complexes from cell lysates, Ab-coated, cross-linked mono-resins were prepared. Protein A Sepharose beads (Abcam, #ab270308) were used to immobilize mouse IgG2a mAbs purified from W6/32 hybridomas (RRID:CVCL_7872) for pan-HLA-I immunopeptidome capture, and L243 hybridomas (RRID:CVCL_4533) for HLA-DR capture. Protein G Sepharose beads (Abcam, #ab270309) were used to immobilize mouse IgG1 mAbs purified from IVA12 hybridomas (RRID:CVCL_G223) for pan-HLA-II immunopeptidome capture. Beads were washed to replace the ethanol storage solution with PBS, then mixed with mAbs at a ratio of 100 µl of actual beads (originally supplied as a 50% agarose slurry in 20% ethanol) per 2 mg mAb, and incubated with gentle rotation for 1 h at 4°C. mAbs were subsequently cross-linked to the beads using 40 mM dimethyl pimelimidate dihydrochloride (DMP; Sigma, #D8388).

IPs were performed according to the CVB3 antigen presentation model used: CaCo2 (W6/32 and IVA12) and DR4/THP-1 cells (L243). Mono-resin mixes were prepared by loading a mAb equivalent of 1 mg per up to 10⁸ cells. mAb-coated beads were incubated with cleared lysates for 16 h at 4°C with gentle rotation. Mono-resins were collected by gravity flow and washed sequentially with 50 mM Tris buffer in four steps: *i)* 150 mM NaCl and 5 mM EDTA, *ii)* 150 mM NaCl, *iii)* 450 mM NaCl, and *iv)* no added salt. Peptides were eluted from the captured pHLA complexes using 10% acetic acid, then passed through a 5-kDa cut-off filter (Millipore, #UFC3LCCNB-HMT), and the flow-through was vacuum-dried. Peptides were resuspended in 1% ACN and 0.1% TFA, desalted on Pierce C18 tips (#84850), and dried by SpeedVac. The final eluates were resuspended in 20 µl of 1% ACN with 0.1% formic acid prior to MS analysis.

Samples were analyzed on a nano-UHPLC system (nanoElute, Bruker Daltonics) coupled to a Trapped Ion Mobility Spectrometry - Time of Flight (timsTOF) SCP (Bruker). Peptides were loaded onto an Aurora C18 column (25 cm × 75 μm, 1.7 μm; IonOpticks) with 0.1% formic acid at 800 bar for 9 min at 50°C. Separation was performed at 50°C using a linear gradient of 2-20% ACN in 0.1% acetic acid over 60 min, then 20-37% ACN in 6 min, at 150 nl/min. Electrospray ionization (CaptiveSpray source, Bruker) settings were 180°C, 1,400 V, and 3 l/min dry gas. MS was acquired in DDA-PASEF mode with one MS survey TIMS-MS and 10 PASEF MS/MS scans per cycle. Ion accumulation and ramp times were 166 ms each, with an ion mobility range of 1/K₀ = 1.7-0.7 Vs cm⁻² and m/z 100-1,700. Multiple-charged and single-charged ions with m/z ≥700 were selected if intensity ≥500 arbitrary units (a.u.), and re-sequenced until a target of 20,000 a.u. was reached. Collision energies for immunopeptidomics were 70 eV at 1/K₀ = 1.7 Vs cm⁻²; 40 eV at 1/K₀ = 1.34 Vs cm⁻²; 40 eV at 1/K₀ = 1.1 Vs cm⁻²; 30 eV at 1/K₀ = 1.06 Vs cm⁻²; and 20 eV at 1/K₀ = 0.7 Vs cm⁻².

TimsTOF data were processed in FragPipe v22.0 (HLA nonspecific workflow) using MSFragger (*69–71*) against a concatenated target-decoy database comprising the UniProtKB/Swiss-Prot human proteome (reviewed entries, downloaded 17/01/2024), and a six-frame translation of the genome sequenced from the CVB3 strain used (<u>Data S1</u>; including all ORFs ≥7 aa); decoys were generated in Philosopher using the reversed sequence method. Searches were run with nonspecific termini, peptide length 7-25 aa, no fixed modifications, and variable modifications including oxidation (M, +15.9949 Da), cysteinylation (C, +119.0041 Da), N-terminal acetylation (+42.0106 Da), pyro-glutamate formation from N-terminal Q (−17.0265 Da) or E (−18.0106 Da), with up to 3 variable modifications per peptide. Precursor and fragment mass tolerances were ±20 ppm. PSMs were rescored with MSBooster (*67*) (DIA-NN models for RT and spectra) and validated with Percolator using a 1% FDR threshold at the PSM, ion, and peptide levels; no protein-level FDR filter was applied. Label-free quantification was performed with IonQuant (*66*) (m/z tolerance 10 ppm, RT tolerance 0.4 min, IM tolerance 0.05 1/K₀, match-between-runs enabled, MaxLFQ normalization). Motif deconvolution were performed using MHCMotifDecon (*72*), and in-silico binding predictions were carried out with NetMHCpan 4.2 (*73*) and NetMHCIIpan 4.3 (*74*). Peptides with a conservative predicted rank ≤4% were classified as binders.

### HLA-A*02:01 binding assays

Peptide binding to HLA-A*02:01 was assessed using a T2 stabilization assay. TAP-deficient T2 cells (HLA-A02:01⁺) were incubated overnight at 26°C, 5% CO_2_ in serum-free RPMI with 10 µM 9-mer peptides (>95% purity; Synpeptide), including Flu MP_58-66_ as a positive control and NY-ESO-1_125-133_ as a negative control. Cells were washed and resuspended in pre-warmed RPMI containing BFA (5 µg/ml) to block new HLA-A2 export. Samples were collected every hour for 5 h, stained with HLA-A2 mAb (RRID:AB_1659245) and Live/Dead Far Red and analyzed on a BD LSRFortessa II flow cytometer. MFI values were normalized to t0 and fitted to a one-phase exponential decay model using GraphPad Prism 8 to calculate half-lives.

For fold-change determination, T2 cells were pulsed overnight at 26°C with peptides at 0.5, 5, or 50 µM in serum-free medium, shifted to 37°C, and treated with 5 µg/ml BFA for 4 h before HLA-A2 staining as above. DMSO-treated cells served as the baseline. Peptides were classified as functional binders if they induced ≥2.3-fold HLA-A2 MFI over the DMSO baseline.

### HLA-DRB1*04:01 binding assays

Peptides predicted to bind HLA-DRB1*04:01 (DR4) (>95% purity; Synpeptide) were assessed using a competitive inhibition assay. DR4/THP-1 cells were pulsed in AIM-V serum-free medium (ThermoFisher, #12055083) with serial dilutions of candidate peptides (0.000001-400 µM) for 2 h at 37°C, followed by addition of the DR4-restricted influenza hemagglutinin (Flu HA)_306-318_ peptide at 0.025 µM for an additional 1 h. This concentration of Flu HA was selected to induce ∼25% of the maximal TNF-α response. After washing, cells were co-cultured in AIM-V medium with a DR4-restricted Flu HA_306-318_-reactive CD4⁺ T cell clone at a 1:5 T cell/THP-1 ratio. BFA (5 µg/ml) was added after 3 h, and co-cultures maintained for a total of 6 h. Cells were then stained with Live/Dead Far Red, fixed/permeabilized, and intracellularly stained with FITC-conjugated anti-TNF-α. Samples were analyzed on a BD BD LSRFortessa II flow cytometer. The percent maximal TNF-α response was plotted against competitor peptide concentration, and curves were fitted in GraphPad Prism 8 using a nonlinear (three parameters) with least squares fitting, constraining bottom=0 and IC_50_>0. Peptides were classified as DR4 binders if they exhibited an IC_50_<400 µM.

### Study participants, HLA typing

Study participants (<u>Table S1</u>) were recruited in Paris after providing written informed consent, under ethics approval 21.01064.000002 (Ouest IV - Nantes). HLA typing was performed by DKMS (Tübingen, Germany), and donors positive for HLA-A*02:01, HLA-DRB1*04:01 or both were selected for T-cell assays. Peripheral blood was collected in 9-mL sodium heparin tubes and PBMCs isolated and cryopreserved following our established protocols (*30, 75*). All experiments were performed on frozen/thawed PBMC samples.

### Combinatorial HLA-I MMr assays

Fluorescent-labeled, peptide loaded HLA-A2 MMrs were generated as previously described (*9, 29–32*), conjugated to fluorochrome-labeled streptavidins at a 1:4 molar ratio, and used at final concentrations of 8-27 nM, adjusted to normalize for the variable staining index of each streptavidin to ensure clear visualization of distinct double-MMr⁺ populations for each fluorochrome pair. PBMCs were thawed in pre-warmed AIM-V medium and rested for 1 h at 37°C with 50 nM dasatinib, followed by negative magnetic enrichment of CD8⁺ T cells (RRID:AB_2728716). Staining with combinatorial, double-coded MMr panels was performed for 30 min at room temperature (RT) in DPBS containing dasatinib, after which phenotyping mAbs were added for 30 min at 4°C: CD8-BUV496 (RRID:AB_2916884), CD45RA-BUV737 (RRID:AB_2870168), PD-1-BB700 (RRID:AB_2744348), CD39-BV605 (RRID:AB_2750430), CCR7-PE/Cy7 (RRID:AB_11126145), CD95-APC/Fire750 (RRID:AB_2629736), TCF-1-AF488 (RRID:AB_2916388) and Live/Dead Aqua (Thermo Fisher, #L34957). After washing, cells were acquired on a Cytek Aurora spectral flow cytometer and analyzed using FlowJo v10 and GraphPad Prism 10. Positive control peptides included MelanA_26–35_ ELA (naïve self-control; ELAGIGILTV; IEDB #12941) and influenza (Flu) MP_58-66_ (recall viral control; GILGFVFTL; IEDB #20354). All MMr⁺ cells were pooled into a concatenated file for t-SNE analysis in FlowJo using all phenotyping panel markers (except the fluorochrome channels assigned to MMrs, CD8, and viability); iterations were set to 1,000, perplexity to 30, and learning rate (eta) to 2,482. The KNN algorithm was set to Exact (Vantage Point Tree), the gradient algorithm to Barnes-Hut, and learning configuration to auto (opt-SNE). Dimensionality reduction was complemented with PhenoGraph clustering (k=30), and results were exported for t-SNE map replotting in Python.

### AIM assays

PBMCs were thawed and rested for 3 h at 37°C, 5% CO_2_ in AIM-V medium before stimulation. 3-4×10⁶ PBMCs were plated in 1 ml AIM-V medium per well in 48-well plates, supplemented with a blocking anti-CD40 mAb (RRID:AB_10839704; 1 µg/ml) to limit CD40L downregulation. Cells were stimulated for 24 h with individual DRB1*04:01-restricted CVB3 peptides (10 µM), DMSO diluent as a negative control, or Flu HA_306-318_ as positive effector/memory control.

Following stimulation, staining was performed in 3 sequential steps separated by DPBS washes in-between. First, cells were incubated at RT for 30 min with chemokine receptor mAbs: CCR4-BV421 (RRID:AB_2737663), CCR6-BUV737 (RRID:AB_2870109), CXCR3-RB705 (RRID:AB_3685840) and CXCR5-BV480 (RRID:AB_2739586). Second, Live/Dead Green viability staining at RT for 20 min (ThermoFisher, #L23101). Third, cells were incubated at 4°C for 30 min with the remaining mAbs to explore T-cell phenotype and activation: CD3-BUV496 (RRID:AB_2870222), CD4-BV805 (RRID:AB_2870176), CD8-BUV563 (RRID:AB_2870199), HLA-DR-BUV661 (RRID:AB_2870252), CD25-BV711 (RRID:AB_2738037), CD127-BV785 (RRID:AB_11219610), CD45RA-BV605 (RRID:AB_11126164), CD69-PE/Dazzle594 (RRID:AB_2564277), CD134 (OX40)-PE/Cy5 (RRID:AB_2894615), CD137 (4-1BB)-PE/Cy7 (RRID:AB_2207741), CD154 (CD40L)-PE (RRID:AB_2751142), and PD-L1-PE/Fire810 (RRID:AB_2894668).

Samples were acquired on a Cytek Aurora spectral flow cytometer and analyzed with FlowJo v10 and GraphPad Prism 10. Boolean gating identified all AIM combinations per donor; combinations absent in matched DMSO stimulation controls were, after redundancy reduction by OR gating, defined as antigen-specific and used for downstream analyses.

### Statistical analysis

Data are shown as median (range) or mean ± SD. Significance was assessed using two-tailed tests with a cutoff value of α = 0.05, as detailed for each figure.

## Acknowledgments

We thank Dr. Hanqing Liao (University of Dundee) for assistance with immunopeptidomics bioinformatics analyses and Dr. Lindsay Whitton (Scripps Research, San Diego) for providing the CVB3-GFP strain. This project has received funding from the European Union’s Horizon Europe research and innovation program ENT1DEP project under Grant Agreement 101137457. Views and opinions expressed are however those of the author(s) only and do not necessarily reflect those of the European Union or of the European Health and Digital Executive Agency (HaDEA). Neither the European Union nor the granting authority can be held responsible for them.

## Author contributions

Conceptualization: OBM, FV, MFT, NT, RM

Methodology: OBM, FV, FS, ZZ, BB, AC, AL, VD, KV, AKL, SY, MFT, NT, RM

Resources: ZZ, BB, AC, VD, KV, AKL, SY, MFT, NT, RM

Investigation: OBM, FV, MP, AL, VD, KV, MFT, NT, RM

Formal analysis: OBM, FV, NT, RM

Visualization: OBM, FV, RM

Supervision: KV, SY, MFT, NT, RM

Funding acquisition: KV, SY, MFT, RM

Project administration: OBM, FV, KV, SY, MFT, RM

Writing—original draft: OBM, RM

Writing—review & editing: OBM, FV, FS, AC, SY, MFT, NT, RM

## Data and materials availability

Immunopeptidomics datasets have been deposited under PRIDE: PXD070471. Proteomics datasets have been deposited under PRIDE: PXD070119. All other data needed to evaluate the conclusions in the paper are available in the main text or the supplementary materials. All unique/stable reagents generated in this study can be provided by R.M. pending scientific review and a completed materials transfer agreement. Requests should be submitted to.

## References

1. C. C. Patterson, V. Harjutsalo, J. Rosenbauer, A. Neu, O. Cinek, T. Skrivarhaug, B. Rami-Merhar, G. Soltesz, J. Svensson, R. C. Parslow, C. Castell, E. J. Schoenle, P. J. Bingley, G. Dahlquist, P. K. Jarosz-Chobot, D. Marciulionyte, E. F. Roche, U. Rothe, N. Bratina, C. Ionescu-Tirgoviste, I. Weets, M. Kocova, V. Cherubini, N. Rojnic Putarek, C. E. deBeaufort, M. Samardzic, A. Green, Trends and cyclical variation in the incidence of childhood type 1 diabetes in 26 European centres in the 25 year period 1989-2013: a multicentre prospective registration study. Diabetologia 62, 408–417 (2019).

2. V. Parikka, K. Nanto-Salonen, M. Saarinen, T. Simell, J. Ilonen, H. Hyoty, R. Veijola, M. Knip, O. Simell, Early seroconversion and rapidly increasing autoantibody concentrations predict prepubertal manifestation of type 1 diabetes in children at genetic risk. Diabetologia 55, 1926–1936 (2012).

3. A. Carré, F. Vecchio, M. Flodstrom-Tullberg, S. You, R. Mallone, Coxsackievirus and type 1 diabetes: diabetogenic mechanisms and implications for prevention. Endocr Rev 44, 737–751 (2023).

4. L. Krogvold, B. Edwin, T. Buanes, G. Frisk, O. Skog, M. Anagandula, O. Korsgren, D. Undlien, M. C. Eike, S. J. Richardson, P. Leete, N. G. Morgan, S. Oikarinen, M. Oikarinen, J. E. Laiho, H. Hyoty, J. Ludvigsson, K. F. Hanssen, K. Dahl-Jorgensen, Detection of a low-grade enteroviral infection in the islets of langerhans of living patients newly diagnosed with type 1 diabetes. Diabetes 64, 1682–1687 (2015).

5. T. Rodriguez-Calvo, J. E. Laiho, M. Oikarinen, P. Akhbari, C. Flaxman, T. Worthington, P. Apaolaza, J. S. Kaddis, I. Kusmartseva, S. Tauriainen, M. Campbell-Thompson, M. A. Atkinson, M. von Herrath, H. Hyöty, N. G. Morgan, A. Pugliese, S. J. Richardson, P. O. D. V. g. for the n, Enterovirus VP1 protein and HLA class I hyperexpression in pancreatic islet cells of organ donors with type 1 diabetes. Diabetologia 68, 1197–1210 (2025).

6. S. J. Richardson, T. Rodriguez-Calvo, J. E. Laiho, J. S. Kaddis, J. O. Nyalwidhe, I. Kusmartseva, S. Morfopoulou, J. F. Petrosino, V. Plagnol, K. Maedler, M. A. Morris, J. L. Nadler, M. A. Atkinson, M. von Herrath, R. E. Lloyd, H. Hyoty, N. G. Morgan, A. Pugliese, P. O. D. V. G. for the n, Joint analysis of the nPOD-Virus Group data: the association of enterovirus with type 1 diabetes is supported by multiple markers of infection in pancreas tissue. Diabetologia 68, 1226–1241 (2025).

7. M. P. Nekoua, E. K. Alidjinou, D. Hober, Persistent coxsackievirus B infection and pathogenesis of type 1 diabetes mellitus. Nat Rev Endocrinol 18, 503–516 (2022).

8. K. Vehik, K. F. Lynch, M. C. Wong, X. Tian, M. C. Ross, R. A. Gibbs, N. J. Ajami, J. F. Petrosino, M. Rewers, J. Toppari, A. G. Ziegler, J. X. She, A. Lernmark, B. Akolkar, W. A. Hagopian, D. A. Schatz, J. P. Krischer, H. Hyoty, R. E. Lloyd, T. S. Group, Prospective virome analyses in young children at increased genetic risk for type 1 diabetes. Nat Med 25, 1865–1872 (2019).

9. F. Vecchio, A. Carre, D. Korenkov, Z. Zhou, P. Apaolaza, S. Tuomela, O. Burgos-Morales, I. Snowhite, J. Perez-Hernandez, B. Brandao, G. Afonso, C. Halliez, J. Kaddis, S. C. Kent, M. Nakayama, S. J. Richardson, J. Vinh, Y. Verdier, J. Laiho, R. Scharfmann, M. Solimena, Z. Marinicova, E. Bismuth, N. Lucidarme, J. Sanchez, C. Bustamante, P. Gomez, S. Buus, P. O. D. V. W. G. n, S. You, A. Pugliese, H. Hyoty, T. Rodriguez-Calvo, M. Flodstrom-Tullberg, R. Mallone, Coxsackievirus infection induces direct pancreatic beta cell killing but poor antiviral CD8(+) T cell responses. Sci Adv 10, eadl1122 (2024).

10. L. Krogvold, I. M. Mynarek, E. Ponzi, F. B. Mørk, T. W. Hessel, T. Roald, N. Lindblom, J. Westman, P. Barker, H. Hyöty, J. Ludvigsson, K. F. Hanssen, J. Johannesen, K. Dahl-Jørgensen, Pleconaril and ribavirin in new-onset type 1 diabetes: a phase 2 randomized trial. Nat Med 29, 2902–2908 (2023).

11. J. L. Dunne, S. J. Richardson, M. A. Atkinson, M. E. Craig, K. Dahl-Jorgensen, M. Flodstrom-Tullberg, H. Hyoty, R. A. Insel, A. Lernmark, R. E. Lloyd, N. G. Morgan, A. Pugliese, Rationale for enteroviral vaccination and antiviral therapies in human type 1 diabetes. Diabetologia 62, 744–753 (2019).

12. V. M. Stone, M. M. Hankaniemi, O. H. Laitinen, A. B. Sioofy-Khojine, A. Lin, I. M. Diaz Lozano, M. A. Mazur, V. Marjomaki, K. Lore, H. Hyoty, V. P. Hytonen, M. Flodstrom-Tullberg, A hexavalent Coxsackievirus B vaccine is highly immunogenic and has a strong protective capacity in mice and nonhuman primates. Sci Adv 6, eaaz2433 (2020).

13. H. Hyoty, S. Kaariainen, J. E. Laiho, G. M. Comer, W. Tian, T. Harkonen, J. P. Lehtonen, S. Oikarinen, L. Puustinen, M. Snyder, F. Leon, M. Scheinin, M. Knip, M. Sanjuan, Safety, tolerability and immunogenicity of PRV-101, a multivalent vaccine targeting coxsackie B viruses (CVBs) associated with type 1 diabetes: a double-blind randomised placebo-controlled Phase I trial. Diabetologia 67, 811–821 (2024).

14. M. P. Ashton, A. Eugster, D. Walther, N. Daehling, S. Riethausen, D. Kuehn, K. Klingel, A. Beyerlein, S. Zillmer, A. G. Ziegler, E. Bonifacio, Incomplete immune response to coxsackie B viruses associates with early autoimmunity against insulin. Sci Rep 6, 32899 (2016).

15. M. Oikarinen, S. Tauriainen, S. Oikarinen, T. Honkanen, P. Collin, I. Rantala, M. Mäki, K. Kaukinen, H. Hyöty, Type 1 Diabetes Is Associated With Enterovirus Infection in Gut Mucosa. Diabetes 61, 687–691 (2012).

16. K. W. Kim, J. L. Horton, C. N. I. Pang, K. Jain, P. Leung, S. R. Isaacs, R. A. Bull, F. Luciani, M. R. Wilkins, J. Catteau, W. I. Lipkin, W. D. Rawlinson, T. Briese, M. E. Craig, Higher abundance of enterovirus A species in the gut of children with islet autoimmunity. Sci Rep 9, 1749 (2019).

17. C. E. Sarraf, C. S. McCormick, G. R. Brown, Y. E. Price, P. A. Hall, D. P. Lane, M. R. Alison, Proliferating cell nuclear antigen immunolocalization in gastro-intestinal epithelia. Digestion 50, 85–91 (1991).

18. V. H. Lugo-Martínez, C. S. Petit, S. Fouquet, J. Le Beyec, J. Chambaz, M. Pinçon-Raymond, P. Cardot, S. Thenet, Epidermal growth factor receptor is involved in enterocyte anoikis through the dismantling of E-cadherin-mediated junctions. Am J Physiol Gastrointest Liver Physiol 296, G235–244 (2009).

19. L. Xiao, J. N. Rao, T. Zou, L. Liu, B. S. Marasa, J. Chen, D. J. Turner, A. Passaniti, J. Y. Wang, Induced JunD in intestinal epithelial cells represses CDK4 transcription through its proximal promoter region following polyamine depletion. Biochem J 403, 573–581 (2007).

20. Z. Wang, Y. Wang, S. Wang, X. Meng, F. Song, W. Huo, S. Zhang, J. Chang, J. Li, B. Zheng, Y. Liu, Y. Zhang, W. Zhang, J. Yu, Coxsackievirus A6 Induces Cell Cycle Arrest in G0/G1 Phase for Viral Production. Front Cell Infect Microbiol 8, 279 (2018).

21. I. V. Sandoval, L. Carrasco, Poliovirus infection and expression of the poliovirus protein 2B provoke the disassembly of the Golgi complex, the organelle target for the antipoliovirus drug Ro-090179. J Virol 71, 4679–4693 (1997).

22. C. A. Quiner, W. T. Jackson, Fragmentation of the Golgi apparatus provides replication membranes for human rhinovirus 1A. Virology 407, 185–195 (2010).

23. M. D. Hansen, I. B. Johnsen, K. A. Stiberg, T. Sherstova, T. Wakita, G. M. Richard, R. K. Kandasamy, E. F. Meurs, M. W. Anthonsen, Hepatitis C virus triggers Golgi fragmentation and autophagy through the immunity-related GTPase M. Proc Natl Acad Sci USA 114, E3462–E3471 (2017).

24. S. Buratta, B. Tancini, K. Sagini, F. Delo, E. Chiaradia, L. Urbanelli, C. Emiliani, Lysosomal Exocytosis, Exosome Release and Secretory Autophagy: The Autophagic- and Endo-Lysosomal Systems Go Extracellular. Int J Mol Sci 21, 2576 (2020).

25. T. A. Solvik, T. A. Nguyen, Y. H. Tony Lin, T. Marsh, E. J. Huang, A. P. Wiita, J. Debnath, A. M. Leidal, Secretory autophagy maintains proteostasis upon lysosome inhibition. J Cell Biol 221, e202110151 (2022).

26. G. V. Shelke, C. D. Williamson, M. Jarnik, J. S. Bonifacino, Inhibition of endolysosome fusion increases exosome secretion. J Cell Biol 222, e202209084 (2023).

27. C. O’Brien, D. R. Flower, C. Feighery, Peptide length significantly influences in vitro affinity for MHC class II molecules. Immunome Res 4, 6 (2008).

28. S. W. Scally, J. Petersen, S. C. Law, N. L. Dudek, H. J. Nel, K. L. Loh, L. C. Wijeyewickrema, S. B. Eckle, J. van Heemst, R. N. Pike, J. McCluskey, R. E. Toes, N. L. La Gruta, A. W. Purcell, H. H. Reid, R. Thomas, J. Rossjohn, A molecular basis for the association of the HLA-DRB1 locus, citrullination, and rheumatoid arthritis. J Exp Med 210, 2569–2582 (2013).

29. S. Culina, A. I. Lalanne, G. Afonso, K. Cerosaletti, S. Pinto, G. Sebastiani, K. Kuranda, L. Nigi, A. Eugster, T. Osterbye, A. Maugein, J. E. McLaren, K. Ladell, E. Larger, J. P. Beressi, A. Lissina, V. Appay, H. W. Davidson, S. Buus, D. A. Price, M. Kuhn, E. Bonifacio, M. Battaglia, S. Caillat-Zucman, F. Dotta, R. Scharfmann, B. Kyewski, R. Mallone, ImMaDiab Study Group, Islet-reactive CD8+ T cell frequencies in the pancreas, but not in blood, distinguish type 1 diabetic patients from healthy donors. Sci Immunol 3, eaao4013 (2018).

30. S. Gonzalez-Duque, M. E. Azoury, M. L. Colli, G. Afonso, J. V. Turatsinze, L. Nigi, A. I. Lalanne, G. Sebastiani, A. Carre, S. Pinto, S. Culina, N. Corcos, M. Bugliani, P. Marchetti, M. Armanet, M. Diedisheim, B. Kyewski, L. M. Steinmetz, S. Buus, S. You, D. Dubois-Laforgue, E. Larger, J. P. Beressi, G. Bruno, F. Dotta, R. Scharfmann, D. L. Eizirik, Y. Verdier, J. Vinh, R. Mallone, Conventional and Neo-antigenic Peptides Presented by beta Cells Are Targeted by Circulating Naive CD8+ T Cells in Type 1 Diabetic and Healthy Donors. Cell Metab 28, 946–960 e946 (2018).

31. M. E. Azoury, M. Tarayrah, G. Afonso, A. Pais, M. L. Colli, C. Maillard, C. Lavaud, L. Alexandre-Heymann, S. Gonzalez-Duque, Y. Verdier, J. Vinh, S. Pinto, S. Buus, D. Dubois-Laforgue, E. Larger, J. P. Beressi, G. Bruno, D. L. Eizirik, S. You, R. Mallone, Peptides Derived From Insulin Granule Proteins are Targeted by CD8+ T Cells Across MHC Class I Restrictions in Humans and NOD Mice. Diabetes 69, 2678–2690 (2020).

32. M. E. Azoury, F. Samassa, M. Buitinga, L. Nigi, N. Brusco, A. Callebaut, M. Giraud, M. Irla, A. I. Lalanne, A. Carre, G. Afonso, Z. Zhou, B. Brandao, M. L. Colli, G. Sebastiani, F. Dotta, M. Nakayama, D. L. Eizirik, S. You, S. Pinto, M. J. Mamula, Y. Verdier, J. Vinh, S. Buus, C. Mathieu, L. Overbergh, R. Mallone, CD8+ T cells variably recognize native versus citrullinated GRP78 epitopes in type 1 diabetes. Diabetes 70, 2879–2891 (2021).

33. C. Poloni, C. Schonhofer, S. Ivison, M. K. Levings, T. S. Steiner, L. Cook, T-cell activation-induced marker assays in health and disease. Immunol Cell Biol 101, 491–503 (2023).

34. O. H. Laitinen, E. Svedin, S. Kapell, A. Nurminen, V. P. Hytonen, M. Flodstrom-Tullberg, Enteroviral proteases: structure, host interactions and pathogenicity. Rev Med Virol 26, 251–267 (2016).

35. A. Mukherjee, S. A. Morosky, E. Delorme-Axford, N. Dybdahl-Sissoko, M. S. Oberste, T. Wang, C. B. Coyne, The coxsackievirus B 3C protease cleaves MAVS and TRIF to attenuate host type I interferon and apoptotic signaling. PLoS Pathog 7, e1001311 (2011).

36. C. T. Cornell, W. B. Kiosses, S. Harkins, J. L. Whitton, Inhibition of protein trafficking by coxsackievirus b3: multiple viral proteins target a single organelle. J Virol 80, 6637–6647 (2006).

37. E. Wessels, D. Duijsings, R. A. Notebaart, W. J. Melchers, F. J. van Kuppeveld, A proline-rich region in the coxsackievirus 3A protein is required for the protein to inhibit endoplasmic reticulum-to-golgi transport. J Virol 79, 5163–5173 (2005).

38. S. B. Deitz, D. A. Dodd, S. Cooper, P. Parham, K. Kirkegaard, MHC I-dependent antigen presentation is inhibited by poliovirus protein 3A. Proc Natl Acad Sci USA 97, 13790–13795 (2000).

39. S. S. Bappy, M. M. Haque Asim, M. M. Ahasan, A. Ahsan, S. Sultana, R. Khanam, A. Z. Shibly, Y. Kabir, Virus-induced host cell metabolic alteration. Rev Med Virol 34, e2505 (2024).

40. S. R. Starck, S. Cardinaud, N. Shastri, Immune surveillance obstructed by viral mRNA. Proc Natl Acad Sci USA 105, 9135–9136 (2008).

41. A. Christiaansen, S. M. Varga, J. V. Spencer, Viral manipulation of the host immune response. Curr Opin Immunol 36, 54–60 (2015).

42. Y. Mohamud, J. Shi, J. Qu, T. Poon, Y. C. Xue, H. Deng, J. Zhang, H. Luo, Enteroviral Infection Inhibits Autophagic Flux via Disruption of the SNARE Complex to Enhance Viral Replication. Cell Rep 22, 3292–3303 (2018).

43. O. Paloheimo, T. O. Ihalainen, S. Tauriainen, O. Välilehto, S. Kirjavainen, E. A. Niskanen, J. P. Laakkonen, H. Hyöty, M. Vihinen-Ranta, Coxsackievirus B3-induced cellular protrusions: structural characteristics and functional competence. J Virol 85, 6714–6724 (2011).

44. S. M. Robinson, G. Tsueng, J. Sin, V. Mangale, S. Rahawi, L. L. McIntyre, W. Williams, N. Kha, C. Cruz, B. M. Hancock, D. P. Nguyen, M. R. Sayen, B. J. Hilton, K. S. Doran, A. M. Segall, R. Wolkowicz, C. T. Cornell, J. L. Whitton, R. A. Gottlieb, R. Feuer, Coxsackievirus B exits the host cell in shed microvesicles displaying autophagosomal markers. PLoS Pathog 10, e1004045 (2014).

45. S. L. Swain, K. K. McKinstry, T. M. Strutt, Expanding roles for CD4(+) T cells in immunity to viruses. Nat Rev Immunol 12, 136–148 (2012).

46. K. H. G. Mills, IL-17 and IL-17-producing cells in protection versus pathology. Nat Rev Immunol 23, 38–54 (2023).

47. D. Fairweather, K. A. Stafford, Y. K. Sung, Update on coxsackievirus B3 myocarditis. Curr Opin Rheumatol 24, 401–407 (2012).

48. I. A. Paiva, J. Badolato-Correa, D. Familiar-Macedo, L. M. de-Oliveira-Pinto, Th17 Cells in Viral Infections-Friend or Foe? Cells 10, 1159 (2021).

49. A. Grifoni, S. Mahajan, J. Sidney, S. Martini, R. H. Scheuermann, B. Peters, A. Sette, A survey of known immune epitopes in the enteroviruses strains associated with acute flaccid myelitis. Hum Immunol 80, 923–929 (2019).

50. R. Varela-Calvino, R. Ellis, G. Sgarbi, C. M. Dayan, M. Peakman, Characterization of the T-cell response to coxsackievirus B4: evidence that effector memory cells predominate in patients with type 1 diabetes. Diabetes 51, 1745–1753 (2002).

51. 51. O. Jensen, S. Trivedi, J. D. Meier, K. C. Fairfax, J. S. Hale, D. T. Leung, A subset of follicular helper-like MAIT cells can provide B cell help and support antibody production in the mucosa. Sci Immunol 7, eabe8931 (2022).

52. T. E. Pankhurst, K. H. Buick, J. L. Lange, A. J. Marshall, K. R. Button, O. R. Palmer, K. J. Farrand, I. Montgomerie, T. W. Bird, N. C. Mason, J. Kuang, B. J. Compton, D. Comoletti, M. Salio, V. Cerundolo, M. E. Quinones-Mateu, G. F. Painter, I. F. Hermans, L. M. Connor, MAIT cells activate dendritic cells to promote T(FH) cell differentiation and induce humoral immunity. Cell Rep 42, 112310 (2023).

53. E. A. Leadbetter, M. C. I. Karlsson, Reading the room: iNKT cells influence B cell responses. Mol Immunol 130, 49–54 (2021).

54. S. Ryu, M. Lim, J. Kim, H. Y. Kim, Versatile roles of innate lymphoid cells at the mucosal barrier: from homeostasis to pathological inflammation. Exp Mol Med 55, 1845–1857 (2023).

55. K. G. Mohn, I. Smith, H. Sjursen, R. J. Cox, Immune responses after live attenuated influenza vaccination. Hum Vaccin Immunother 14, 571–578 (2018).

56. M. T. Yeh, M. Smith, S. Carlyle, J. L. Konopka-Anstadt, C. C. Burns, J. Konz, R. Andino, A. Macadam, Genetic stabilization of attenuated oral vaccines against poliovirus types 1 and 3. Nature 619, 135–142 (2023).

57. L. Rivera Mejía, L. Peña Méndez, A. S. Bandyopadhyay, C. Gast, S. Mazara, K. Rodriguez, N. Rosario, Y. Zhang, B. A. Mainou, J. Jimeno, G. Aguirre, R. Rüttimann, Safety and immunogenicity of shorter interval schedules of the novel oral poliovirus vaccine type 2 in infants: a phase 3, randomised, controlled, non-inferiority study in the Dominican Republic. Lancet Infect Dis, 24, 275–284 (2023).

58. L. Bauer, R. Manganaro, B. Zonsics, J. Strating, P. El Kazzi, M. Lorenzo Lopez, R. Ulferts, C. van Hoey, M. J. Mate, T. Langer, B. Coutard, A. Brancale, F. J. M. van Kuppeveld, Fluoxetine Inhibits Enterovirus Replication by Targeting the Viral 2C Protein in a Stereospecific Manner. ACS Infect Dis 5, 1609–1623 (2019).

59. C. C. Kemball, S. Harkins, J. K. Whitmire, C. T. Flynn, R. Feuer, J. L. Whitton, Coxsackievirus B3 inhibits antigen presentation in vivo, exerting a profound and selective effect on the MHC class I pathway. PLoS Pathog 5, e1000618 (2009).

60. Y. Mohamud, J. C. Lin, S. W. Hwang, A. Bahreyni, Z. C. Wang, H. Luo, Coxsackievirus B3 Activates Macrophages Independently of CAR-Mediated Viral Entry. Viruses 16, 1456 (2024).

61. R. Feuer, I. Mena, R. Pagarigan, M. K. Slifka, J. L. Whitton, Cell cycle status affects coxsackievirus replication, persistence, and reactivation in vitro. J Virol 76, 4430–4440 (2002).

62. N. V. V. Saarinen, J. E. Laiho, S. J. Richardson, M. Zeissler, V. M. Stone, V. Marjomäki, T. Kantoluoto, M. S. Horwitz, A. Sioofy-Khojine, A. Honkimaa, A novel rat CVB1-VP1 monoclonal antibody 3A6 detects a broad range of enteroviruses. Sci Rep 8, 1–13 (2018).

63. A. Carré, F. Samassa, Z. Zhou, J. Perez-Hernandez, C. Lekka, A. Manganaro, M. Oshima, H. Liao, R. Parker, A. Nicastri, B. Brandao, M. L. Colli, D. L. Eizirik, J. Aluri, D. Patel, M. Göransson, O. Burgos Morales, A. Anderson, L. Landry, F. Kobaisi, R. Scharfmann, L. Marselli, P. Marchetti, S. You, M. Nakayama, S. R. Hadrup, S. C. Kent, S. J. Richardson, N. Ternette, R. Mallone, Interferon-α promotes HLA-B-restricted presentation of conventional and alternative antigens in human pancreatic β-cells. Nat Commun 16, 765 (2025).

64. M. Genin, F. Clement, A. Fattaccioli, M. Raes, C. Michiels, M1 and M2 macrophages derived from THP-1 cells differentially modulate the response of cancer cells to etoposide. BMC Cancer 15, 577 (2015).

65. M. Daigneault, J. A. Preston, H. M. Marriott, M. K. Whyte, D. H. Dockrell, The identification of markers of macrophage differentiation in PMA-stimulated THP-1 cells and monocyte-derived macrophages. PLoS One 5, e8668 (2010).

66. F. Yu, S. E. Haynes, A. I. Nesvizhskii, IonQuant Enables Accurate and Sensitive Label-Free Quantification With FDR-Controlled Match-Between-Runs. Mol Cell Proteomics 20, 100077 (2021).

67. K. L. Yang, F. Yu, G. C. Teo, K. Li, V. Demichev, M. Ralser, A. I. Nesvizhskii, MSBooster: improving peptide identification rates using deep learning-based features. Nat Commun 14, 4539 (2023).

68. Y. Hsiao, H. Zhang, G. X. Li, Y. Deng, F. Yu, H. Valipour Kahrood, J. R. Steele, R. B. Schittenhelm, A. I. Nesvizhskii, Analysis and Visualization of Quantitative Proteomics Data Using FragPipe-Analyst. J Proteome Res 23, 4303–4315 (2024).

69. A. T. Kong, F. V. Leprevost, D. M. Avtonomov, D. Mellacheruvu, A. I. Nesvizhskii, MSFragger: ultrafast and comprehensive peptide identification in mass spectrometry-based proteomics. Nat Methods 14, 513–520 (2017).

70. F. Yu, S. E. Haynes, G. C. Teo, D. M. Avtonomov, D. A. Polasky, A. I. Nesvizhskii, Fast Quantitative Analysis of timsTOF PASEF Data with MSFragger and IonQuant. Mol Cell Proteomics 19, 1575–1585 (2020).

71. G. C. Teo, D. A. Polasky, F. Yu, A. I. Nesvizhskii, Fast Deisotoping Algorithm and Its Implementation in the MSFragger Search Engine. J Proteome Res 20, 498–505 (2021).

72. S. Kaabinejadian, C. Barra, B. Alvarez, H. Yari, W. H. Hildebrand, M. Nielsen, Accurate MHC Motif Deconvolution of Immunopeptidomics Data Reveals a Significant Contribution of DRB3, 4 and 5 to the Total DR Immunopeptidome. Front Immunol 13, 835454 (2022).

73. J. B. Nilsson, J. Greenbaum, B. Peters, M. Nielsen, NetMHCpan-4.2: improved prediction of CD8+ epitopes by use of transfer learning and structural features. Front Immunol 16, 1616113 (2025).

74. J. B. Nilsson, S. Kaabinejadian, H. Yari, M. G. D. Kester, P. van Balen, W. H. Hildebrand, M. Nielsen, Accurate prediction of HLA class II antigen presentation across all loci using tailored data acquisition and refined machine learning. Sci Adv 9, eadj6367 (2023).

75. R. Mallone, S. I. Mannering, B. M. Brooks-Worrell, I. Durinovic-Bello, C. M. Cilio, F. S. Wong, N. C. Schloot, Isolation and preservation of peripheral blood mononuclear cells for analysis of islet antigen-reactive T cell responses: position statement of the T-Cell Workshop Committee of the Immunology of Diabetes Society. Clin Exp Immunol 163, 33–49 (2011).

