## Supplementary material for "Coxsackievirus B Escapes Antiviral CD8⁺ T Cells but Triggers Robust CD4⁺ Memory Responses": Fig. S

Orlando Burgos-Morales *et al.*

**This PDF file includes:**

Figs. S1 to S5
Tables S1
Data S1

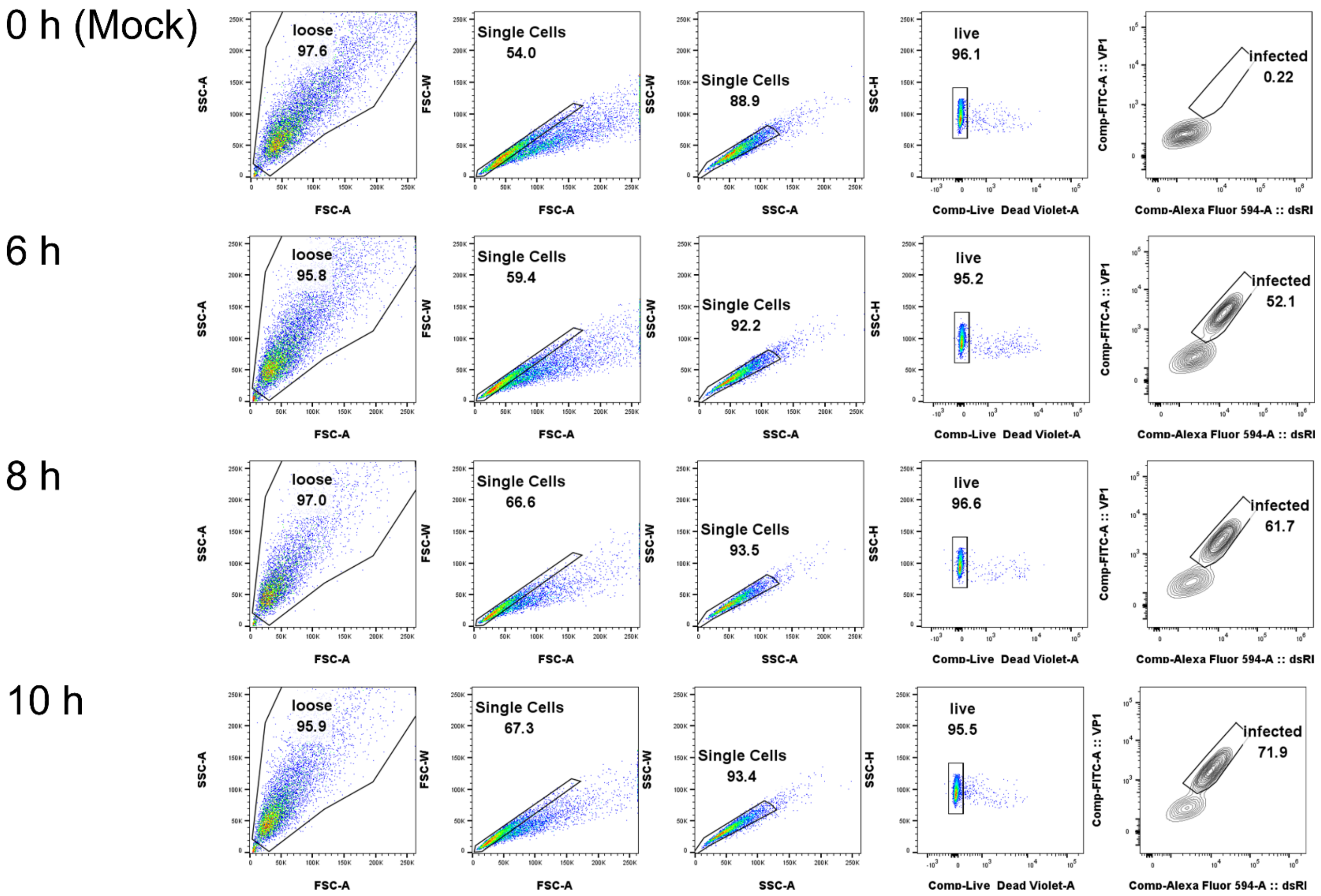


**Fig. S1. Gating strategy used for CVB3 infection time course.** CaCo2 enterocytes ($0.2 \times 10^6$ cells/well, 24-well plate, 70% confluency) were infected at 300 MOI for the indicated time. Cumulative data is shown in Fig. 1A.

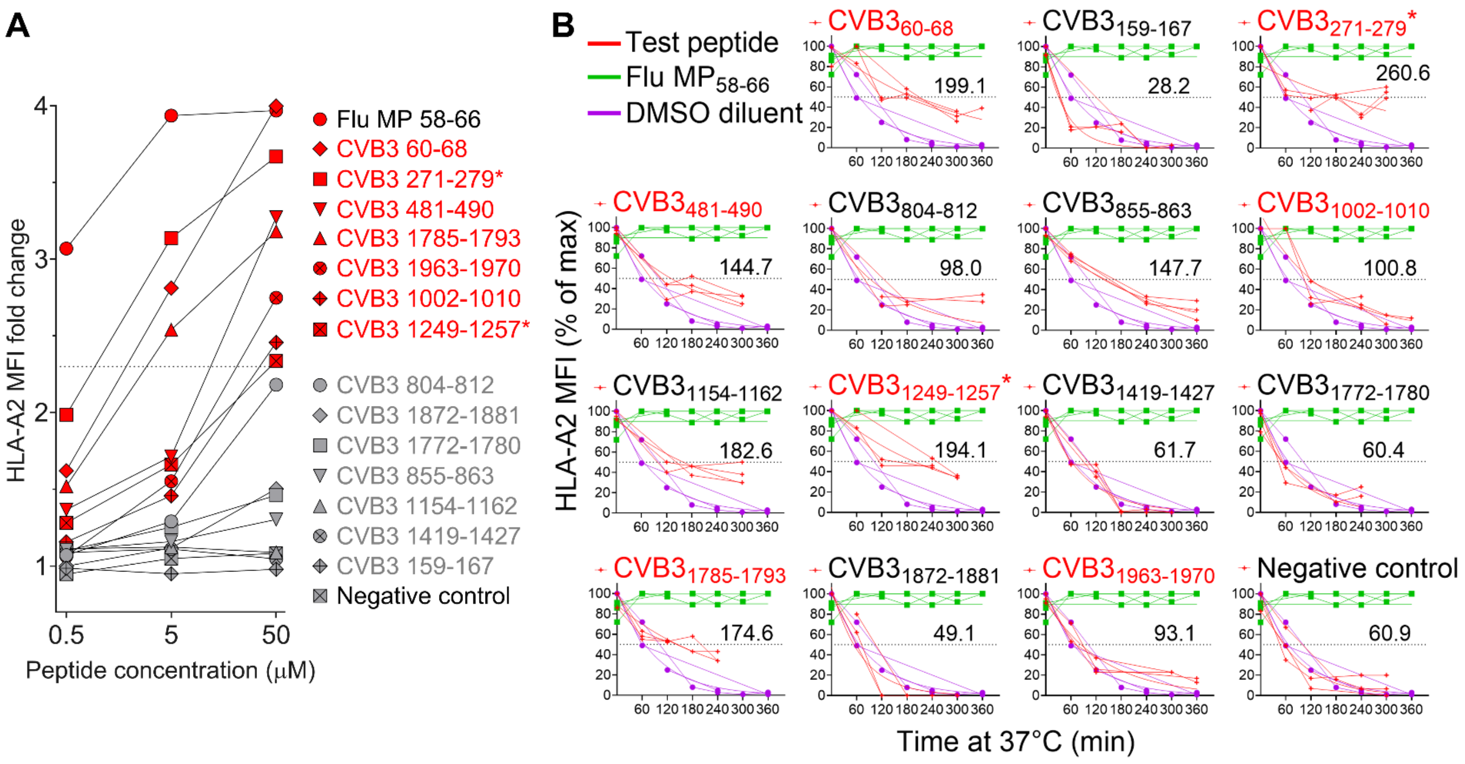


**Fig. S2. HLA-A*02:01 binding measured by T2 HLA-A2 stabilization assays. A.** Peptide binding $K_{on}$, measured as HLA-A2 median fluorescence intensity (MFI) fold change compared with a negative control non-binding peptide (NY-ESO-$1_{125\text{-}133}$) at the indicated peptide concentrations. A strong Flu $MP_{58\text{-}66}$ binder was included as positive control. Peptide $CVB3_{1154\text{-}1162}$ (NetMHCpan rank 8.49%) was included as a negative control. Confirmed HLA-A2 binders (MFI fold change ≥2.3 at 50 µM) are shown in red; peptides below the positive threshold are shown in grey. The $CVB3_{271\text{-}279}$ and $CVB3_{1249\text{-}1257}$ epitopes previously identified in β-cell immunopeptidomes and validated for T-cell recognition are indicated by asterisks. Results refer to a representative experiment out of 3 performed. **B.** Peptide binding $K_{off}$, measured as HLA-A2 MFI decay over time upon incubation at 37°C in the presence of brefeldin A. Results show 3 replicate measurements from a representative experiment out of 2 performed.

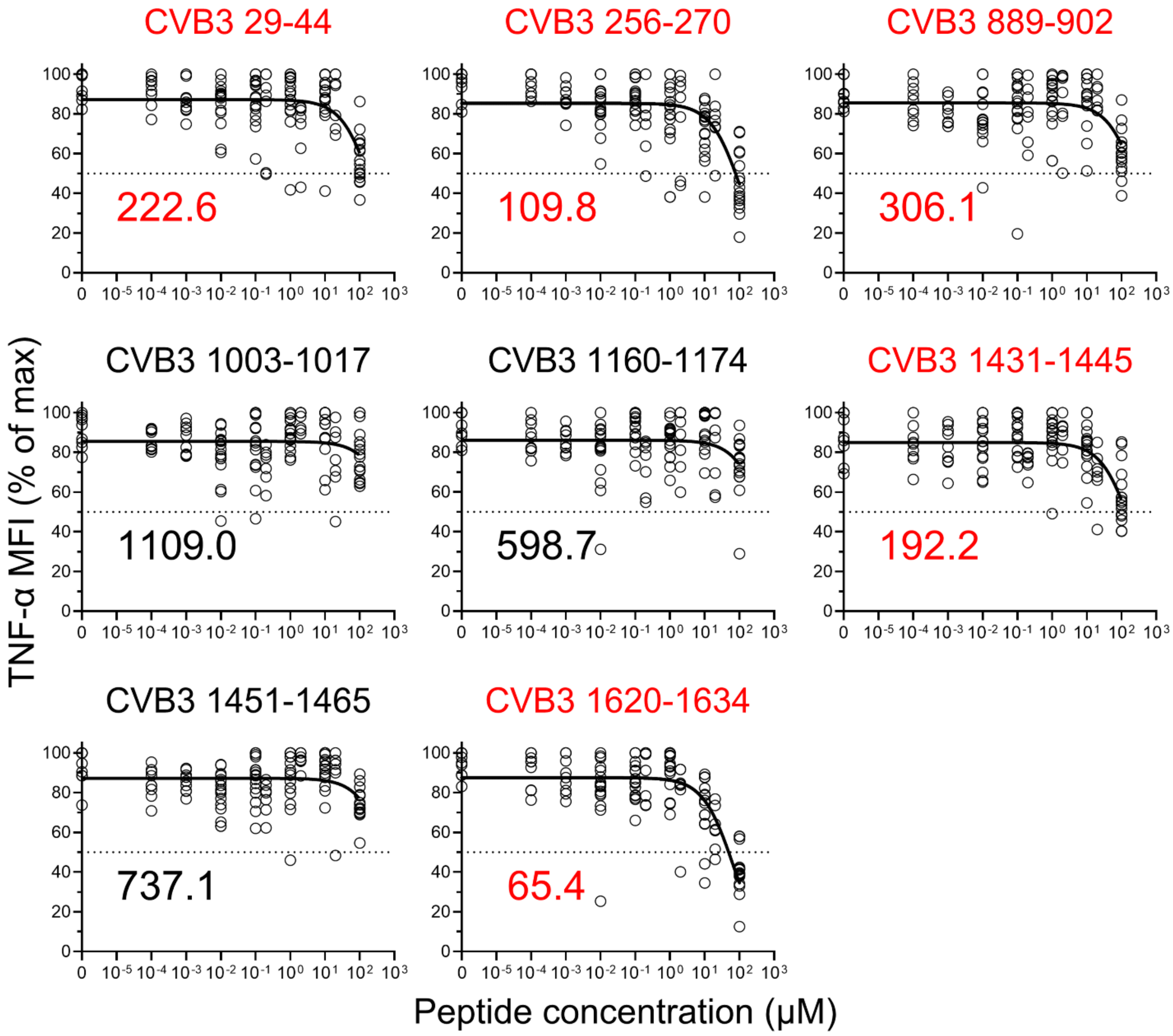


**Fig. S3. HLA-DRB1*04:01 binding measured by a T-cell reporter competition assay.** A DRB1*04:01-restricted Flu $HA_{306\text{-}318}$-reactive $CD4^+$ T-cell clone was used as reporter. DR4/THP-1 cells were pulsed with serial dilutions of test CVB3 peptides, followed by the addition of the cognate Flu $HA_{306\text{-}318}$ peptide. Peptides $CVB3_{1003\text{-}1017}$ and $CVB3_{1160\text{-}1174}$ (NetMHCIIpan rank 7.49% and 6.07%, respectively) were included as negative controls. After washing, T cells were added for 6 h, followed by intracellular TNF-α staining. Confirmed HLA-DRB1*04:01 binders (shown in red) were defined based on a half-maximal inhibitory concentration (IC50) <400 μM of the TNF-α median fluorescence intensity (MFI), noted for each graph. Results show 16 replicates from a representative experiment out of 3 performed.

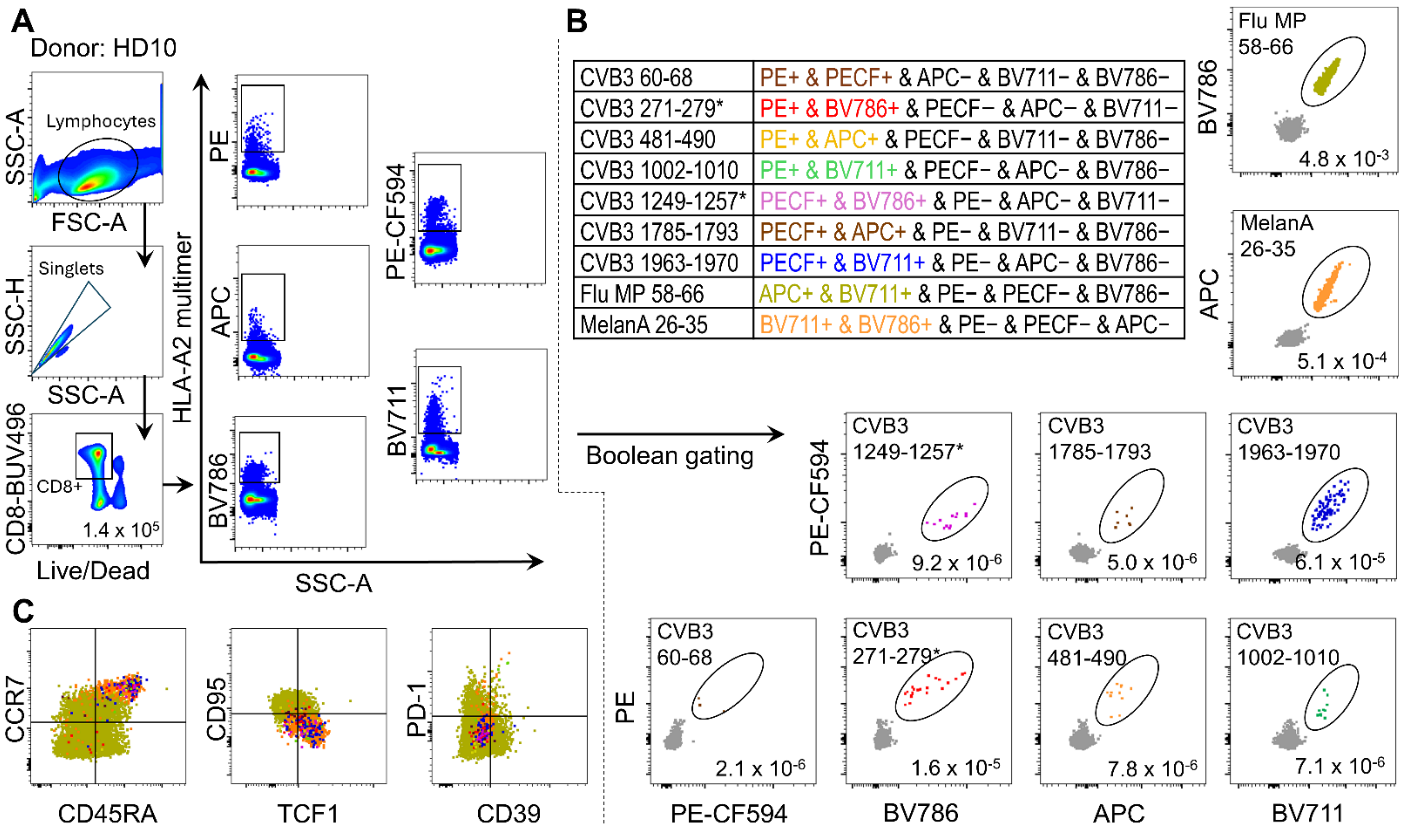


**Fig. S4. Identification of CVB3 peptide-reactive CD8$^+$ T cells using combinatorial HLA-A2 multimer (MMr) assays. A.** Following sequential lymphocyte, singlet and live CD8$^+$ cell gating (*left*), total PE$^+$, PE-CF594$^+$, APC$^+$, BV711$^+$ and BV786$^+$ MMr$^+$ cells were visualized (*right*). **B.** Since each peptide-loaded MMr is assembled with a unique pair of fluorochrome-labeled streptavidins, Boolean operators allowed to selective visualize each double-MMr$^+$ population by including only those events positive for the corresponding fluorochrome pair and excluding all other fluorochromes. The corresponding Boolean gating strategy is listed in the table for each of the 9 peptides analyzed: 7 CVB3 peptides, a Flu MP$_{58-66}$ and a MelanA$_{26-35}$ epitope used as controls for effector/memory and naïve T-cell phenotype, respectively. Dot plots display representative HLA-A2 MMr readout. Events corresponding to each double-MMr$^+$ peptide-reactive T-cell population are shown in color (corresponding to the color code of the table). Numbers in each panel indicate the MMr$^+$CD8$^+$ T-cell frequency out of total CD8$^+$ T cells. The CVB3$_{271-279}$ and CVB3$_{1249-1257}$ epitopes previously identified in β-cell immunopeptidomes and validated for T-cell recognition are indicated by asterisks. **C.** Representative phenotype staining of CVB3 MMr$^+$ CD8$^+$ T cells. All double-MMr$^+$ CD8$^+$ T cells are overlaid in the same dot plots, with each peptide reactivity colored with the corresponding color code of the table.

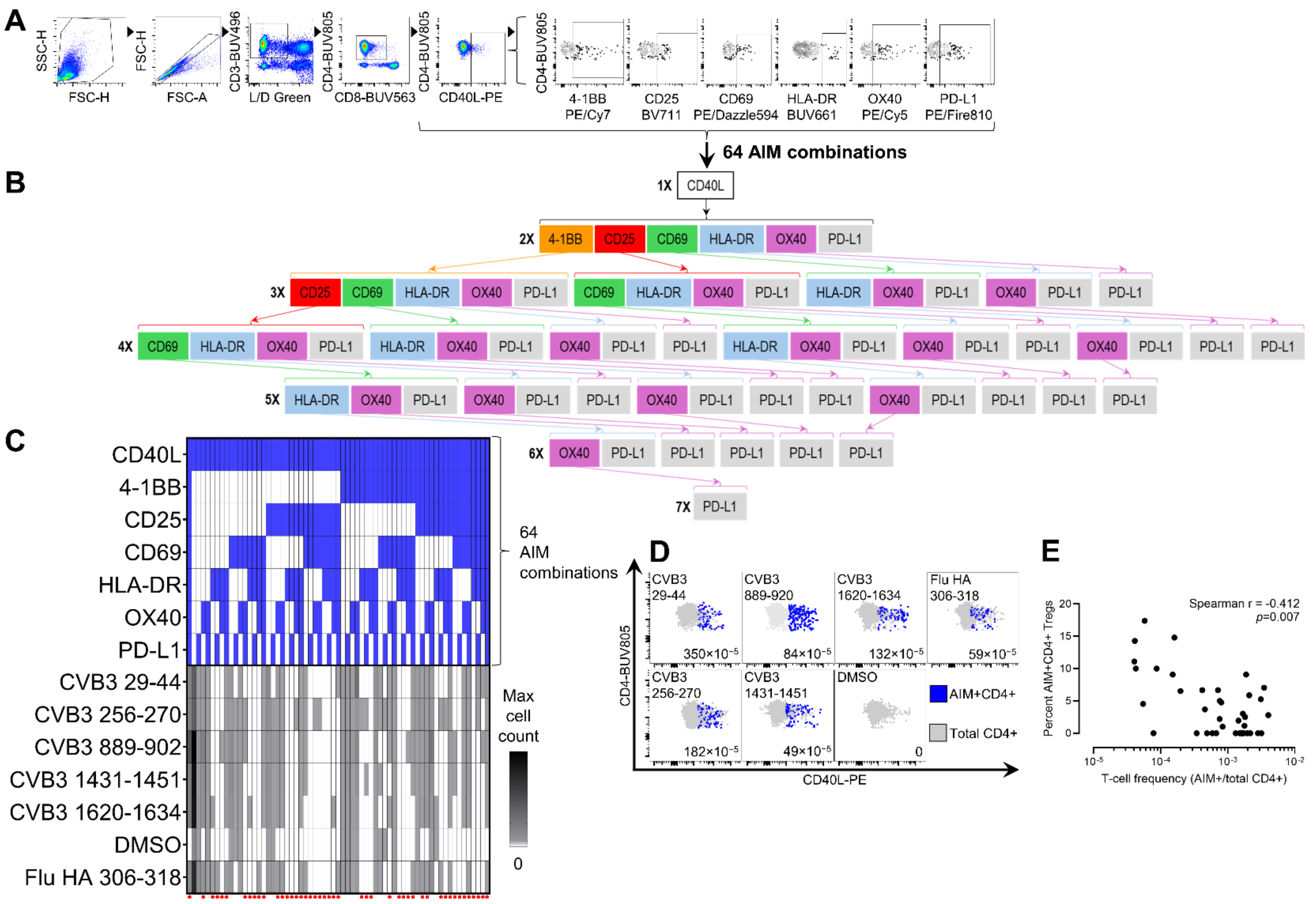


**Fig. S5. Identification of HLA-DRB1*04:01-restricted CVB3 peptide-reactive CD4$^+$ T cells by combinatorial AIM assays.** PBMCs were stimulated with individual peptides for 24 h before flow cytometry. **A.** To detect peptide-reactive AIM$^+$CD4$^+$ T cells, a sequential gating strategy was first applied to identify live, single CD3$^+$CD4$^+$CD8$^-$ T cells, followed by a first gate on CD40L$^+$ cells which was then combined with the other 6 AIMs (4-1BB, CD25, CD69, HLA-DR, OX40, and PD-L1). **B.** All possible 64 AIM combinations with CD40L were analyzed: 1X, CD40L alone; 2X, 1 other AIM; 3X, 2 other AIMs; up to 7X, all 7 AIMs combined. **C.** For each donor, the 64 AIM combinations (*top panel*) were first analyzed to select only those scoring 0 background events in the DMSO negative control stimulation condition (*bottom panel*; 0 counts highlighted in white; combinations marked with a red dot). For each individual peptide stimulation, positive AIM combinations were then retained among those previously selected (*bottom panel*; counts >0 highlighted in grey shades) and further combined with an OR gate to define AIM$^+$CD4$^+$ T cells with maximal specificity and sensitivity. **D.** Representative dot plots displaying the response to the indicated CVB3 peptides, DMSO diluent negative control and the recall Flu HA$_{306\text{-}318}$ epitope. AIM$^+$ and total CD4$^+$ T cells are displayed in blue and grey, respectively; the corresponding frequencies out of total CD4$^+$ T cells are indicated for each dot plot. No AIM$^+$ events were detected in the DMSO condition, validating the stringency of the selection process. **E.** Inverse correlation between AIM$^+$CD4$^+$ T-cell frequencies and percent AIM$^+$CD4$^+$ Tregs recognizing the same CVB3 peptide.

| Code | Age(yrs) | Gender (M/F) | HLA | Symbol |
|---|---|---|---|---|
| HD01 | 51 | M | A2 | ● |
| HD02 | 41 | M | A2/DR4 | ● (grey) |
| HD03 | 32 | F | A2/DR4 | ■ (grey) |
| HD04 | 26 | F | A2 | ■ |
| HD05 | 38 | F | A2 | ▲ (grey) |
| HD06 | 35 | M | DR4 | ▽ |
| HD07 | 32 | M | A2 | ▼ (grey) |
| HD08 | 30 | F | A2 | ◆ |
| HD09 | 35 | F | A2/DR4 | ▲ |
| HD10 | 25 | F | A2/DR4 | □ |
| HD11 | 27 | M | DR4 | ○ |
| HD12 | 25 | F | DR4 | ▼ |

**Table S1. Healthy donors recruited for T-cell experiments.** All HLA genotypes were 4-digit A*02:01 and/or DRB1*04:01.

**Data S1. CVB3 genome sequenced from the strain used in all experiments and the corresponding amino acid translation of the canonical reading frame.**

**>CVB3 genome**

```
TTAAAACAGCCTGTGGGTTGATCCCACCCACAGGGCCCATTGGGCGCTAGCACTCTGGTATCACGGTACCTTTG
TGTGCCTGTTTTATACCCCCTCCCCCAACTGTAACTTAGAAGTAACACACACCGATCAACAGTCAGCGTGGCAC
ACCAGCCACGTTTTGATCAAGCACTTCTGTTACCCCGGACTGAGTATCAATAGACTGCTCACGCGGTTGAAGGA
GAAAGCGTTCGTTATCCGGCCAACTACTTCGAAAAACCTAGTAACACCGTGGAAGTTGCAGAGTGTTTCGCTCA
GCACTACCCCAGTGTAGATCAGGTCGATGAGTCACCGCATTCCCCACGGGCGACCGTGGCGGTGGCTGCGTTGG
CGGCCTGCCCATGGGGAAACCTATGGGACGCTCYAATACAGACATGGTGCGAAGAGTCTATTGAGCTAGTTGGT
AGTCCTCCGGCCCCTGAATGCGGCTAATCCTAACTGCGGAGCACACACCCTCAAGCCAGAGGGCAGTGTGTCGT
AACGGGCAACTCTGCAGCGGAACCGACTACTTTGGGTGTCCGTGTTTCACTTTTATTCCTAYACTGGCTGCTTA
TGGTGACAATTGAGAGATTGTTACCATATAGCTATTGGATTGGCCATCCGGTGACCAATAGAGCTATTATATAT
CTCTTTGTTGGGTTTATACCACTTAGCTTGAAAGAGGTTAAAACATTACAATTCATTGTTAAGTTGAATACAGC
AAAATGGGAGCTCAAGTATCAACGCAAAAGACTGGGGCACATGAGACCGGGCTGAATGCTAGCGGCAATTCCAT
CATTCACTACACAAATATTAATTATTACAAGGATGCCGCATCCAACTCAGCCAATCGGCAGGATTTCACTCAAG
ACCCGGGCAAGTTCACAGAACCAGTAAAAGATATCATGATTAAATCACTACCAGCTCTCAACTCCCCCACAGTA
GAGGAGTGCGGATACAGTGACAGGGCGAGATCAATCACATTAGGTAACTCCACCATAACGACTCAGGAATGCGC
CAACGTGGTGGTGGGCTATGGAGTATGGCCAGATTATCTAAAGGATAGTGAGGCAACAGCAGAGGACCAACCGA
CCCAACCAGACGTTGCCACATGTAGGTTCTATACCCTTGACTCTGTGCAATGGCAGAAAACCTCACCAGGATGG
TGGTGGAAGCTGCCCGATGCTTTGTCGAACTTAGGACTGTTTGGGCAGAACATGCAGTACCACTACTTAGGCCG
AACTGGGTATACCGTACATGTGCAGTGCAATGCATCTAAGTTCCACCAAGGATGCTTGCTAGTAGTGTGTGTAC
CGGAAGCTGAGATGGGTTGCGCAACGCTAGACAACACCCCATCCAGTGCAGAATTGCTGGGGGGCGATAGCGCA
AAAGAGTTTGCGGACAAACCGGTCGCATCCGGGTCCAACAAGTTGGTACAGAGGGTGGTGTATAATGCAGGCAT
GGGGGTGGGTGTTGGAAACCTCACCATTTTCCCCCACCAATGGATCAACCTACGCACCAATAATAGTGCTACAA
TTGTGATGCCATACACCAACAGTGTACCTATGGATAACATGTTTAGGCATAACAACGTCACCCTAATGGTTATC
CCATTTGTACCGCTAGATTACTGCCCTGGGTCCACCACGTACGTCCCAATTACGGTCACGATAGCCCCAATGTG
TGCCGAGTACAATGGGTTACGTTTAGCAGGGCACCAGGGCTTACCAACCATGAATACTCCGGGGAGCTGTCAAT
TTCTGACATCAGACGACTTCCAATCACCATCCGCCATGCCGCAATATGACGTCACACCAGAGATGAGGATACCT
GGTGAGGTGAAGAACTTGATGGAAATAGCTGAGGTTGACTCAGTTGTCCCAGTCCAAAATGTTGGAGAGAAGGT
CAACTCTATGGAAGCATACCAGATACCTGTGAGATCCAATGAAGGATCTGGAACGCAAGTATTCGGCTTTCCAC
TGCAACCAGGGTACTCGAGTGTTTTTAGTCGGACGCTCCTAGGAGAGATCTTGAACTATTATACACATTGGTCA
GGCAGCATAAAGCTTACGTTTATGTTCTGTGGTTCGGCCATGGCTACTGGAAAATTCCTTTTGGCATACTCACC
ACCAGGTGCTGGAGCTCCYACAAAAAGGGTTGATGCCATGCTTGGTACTCATGTAATTTGGGACGTGGGGCTAC
AATCAAGTTGCGTGCTGTGTATACCCTGGATAAGCCAAACACACTACCGGTATGTTGCTTCAGATGAGTATACC
GCAGGGGGTTTYATTACGTGCTGGTATCAAACAAACATAGTGGTCCCAGCGGATGCCCAAAGCTCCTGTTACAT
CATGTGTTTCGTGTCAGCATGCAATGACTTCTCTGTCAGGCTATTGAAGGACACTCCTTTCATTTCGCAGGAAA
ACTTTTTCCAGGGCCCAGTGGAAGACGCGATAACAGCCGCTATAGGGAGAGTTGCGGATACCGTGGGTACAGGG
CCAACCAACTCAGAAGCTATACCAGCACTCACTGCTGCTGAGACAGGTCACACGTCACAAGTAGTGCCGGGTGA
CACCATGCAGACACGCCACGTTAAGAACTACCATTCAAGGTCCGAGTCAACCATAGAGAACTTCCTATGTAGGT
CAGCATGCGTGTACTTTACGGAGTATGAAAACTCAGGTGCCAAGCGGTATGCTGAATGGGTATTAACACCACGA
CAAGCAGCACAACTTAGGAGAAAGCTAGAATTCTTTACCTACGTCCGGTTCGACCTGGAGCTGACGTTTGTCAT
AACAAGTACTCAACAGCCCTCAACCACACAGAACCAAGACGCACAGATCCTAACACACCAAATTATGTATGTAC
CACCAGGTGGACCTGTACCAGATAAAGTTGATTCATACGTGTGGCAAACATCTACGAATCCCAGTGTGTTTTGG
ACCGAGGGAAACGCCCCGCCGCGCATGTCCATACCGTTTTTGAGCATTGGCAACGCCTATTCAAATTTCTATGA
CGGATGGTCTGAATTTTCCAGGAACGGAGTTTACGGCATCAACACGCTAAACAACATGGGCACGCTATATGCAA
GACATGTCAACGCTGGAAGCACGGGTCCAATAAAAAGCACCATTAGAATCTACTTCAAACCGAAGCATGTCAAA
GCGTGGATACCTAGACCACCTAGACTCTGCCAATACGAGAAGGCAAAGAACGTGAACTTCCAACCCAGCGGAGT
TACCACTACTAGGCAAAGCATCACTACAATGACAAATACGGGCGCATTTGGACAACAATCAGGGGCAGCGTATG
TGGGRAACTACAGGGTAGTAAATAGACAYYTAGCTACCAGTGCTGACTGGCAAAACTGTGTGTGGGAAAGTTAC
AACAGAGACCTCTTAGTGAGCACGACCACAGCACATGGATGTGATATTATAGCCAGATGTCAGTGCACAACGGG
AGTGTACTTTTGTGCGTCCAAAAACAAGCACTACCCAATTTCGTTTGAAGGACCAGRTCTAGTAGAGGTCCAAG
AGAGTGAATACTACCCCAGGAGATACCAATCCCATGTGCTTTTAGCAGCTGGATTTTCCGAACCAGGTGACTGT
```

GGCGGTATCCTAAGGTGTGAGCATGGTGTCATTGGCATTGTGACCATGGGGGGTGAAGGCGTGGTCGGCTTTGC
AGACATCCGTGATCTCCTGTGGCTGGAAGATGATGCAATGGAACAGGGAGTGAAGGACTATGTGGAACAGCTTG
GAAATGCATTCGGCTCCGGCTTTACTAACCAAATATGTGAGCAAGTCAACCTCCTGAAAGAATCACTAGTGGGT
CAAGACTCCATCTTAGAGAAATCTCTAAAAGCCTTAGTTAAGATAATATCAGCCTTAGTAATTGTGGTGAGGAA
CCACGATGACCTGATCACTGTGACTGCCACACTAGCCCTTATCGGTTGTACCTCGTCCCCGTGGCGGTGGCTCA
AACAGAAGGKGTCACAATATTACGGAATCCCTATGGCTGAACGCCAAAACAATAGCTGGCTTAAGAAATTTACT
GAAATGACGAATGCTTGCAAGGGTATGGAATGGATAGCTGTCAAAATTCAGAAATTCATTGAATGGCTCAAAGT
AAAAATTTTGCCAGAGGTCAGGGAAAAACACGAATTCCTGAACAGACTTAAACAACTCCCCTTATTAGAAAGTC
AGATCGCCACAATCGAGCAGAGCGCGCCATCCCAAAGTGACCAGGAACAATTATTTTCCAATGTCCAATACTTT
GCCCACTATTGCAGAAAGTACGCTCCCCTCTACGCAGCTGAAGCAAAGAGGGTGTTCTCCCTTGAGAAGAAGAT
GAGCAATTACATACAGTTCAAGTCCAAATGCCGTATTGAACCTGTATGTTTGCTCCTGCACGGGAGCCCTGGTG
CCGGCAAGTCGGTGGCAACAAACTTAATTGGAAGGTCGCTTGCTGAGAAACTCAACAGCTCAGTGTACTCACTA
CCGCCAGACCCAGATCACTTCGACGGATACAAACAGCAGGCCGTGGTGATTATGGACGATCTATGCCAGAATCC
TGATGGGAAAGACGTCTCCTTGTTCTGCCAAATGGTTTCCAGTGTAGATTTTGTACCACCCATGGCTGCCCTAG
AAGAGAAAGGCATTCTGTTCACCTCACCGTTTGTCTTGGCATCGACCAATGCAGGATCTATTAATGCTCCAACC
GTGTCAGATAGCAGAGCCTTGGCAAGGAGATTTCACTTTGACATGAACATCGAGGTTATTTCCATGTACAGTCA
GAATGGCAAGATAAACATGCCCATGTCAGTCAAGACTTGTGACGATGAGTGTTGCCCGGTCAATTTTAAAAAGT
GCTGCCCTCTTGTGTGTGGGAAGGCTATACAATTCATTGATAGAAGAACACAGGTCAGATACTCTCTAGACATG
CTAGTCACCGAGATGTTTAGGGAGTACAATCATAGACATAGCGTGGGGACCACGCTTGAGGCACTGTTCCAGGG
ACCACCAGTATACAGAGAGATCAAAATTAGCGTTGCACCAGAGACACCACCACCGCCCGCCATTGCGGACCTGC
TCAAATCGGTAGACAGTGAGGCTGTGAGGGAGTACTGCAAAGAAAAAGGATGGTTGGTTCCTGAGATCAACTCC
ACCCTCCAAATTGAGAAACATGTCAGTCGGGCTTTCATTTGCTTACAGGCATTGACCACATTTGTGTCAGTGGC
TGGAATCATATATATAATATATAAGCTCTTTGCGGGTTTTCAAGGYGCTTATACAGGAGTGCCCAACCAGAAGC
CCAGAGTGCCTACCCTGAGGCAAGCAAAAGTGCAAGGCCCTGCCTTTGAGTTCGCCGTCGCAATGATGAAAAGG
AACTCAAGCACGGTGAAAACTGAATATGGCGAGTTTACCATGCTGGGCATCTATGACAGGTGGGCCGTTTTGCC
ACGCCACGCCAAACCTGGGCCAACCATCTTGATGAATGATCAAGAGGTTGGTGTGCTAGATGCCAAGGAGCTAG
TAGACAAGGACGGCACCAACTTAGAACTGACACTACTCAAATTGAACCGGAATGAGAAGTTCAGAGACATCAGA
GGCTTCTTAGCCAAGGAGGAAGTGGAGGTTAATGAGGCAGTGCTAGCAATTAACACCAGCAAGTTTCCCAACAT
GTACATTCCAGTAGGACAGGTCACAGAATACGGCTTCCTAAACCTAGGTGGCACACCCACCAAGAGAATGCTTA
TGTACAACTTCCCCACAAGAGCAGGCCAGTGTGGTGGAGTGCTCATGTCCACCGGCAAGGTACTGGGTATCCAT
GTTGGTGGAAATGGCCATCAGGGCTTCTCAGCAGCACTCCTCAAACACTACTTCAATGATGAGCAAGGTGAAAT
AGAATTTATTGAGAGCTCAAAGGACGCCGGGTTTCCAGTCATCAACACACCAAGTAAAACAAAGTTGGAGCCTA
GTGTTTTCCACCAGGTCTTTGAGGGGAACAAAGAACCAGCAGTACTCAGGAGTGGGGATCCTCGTCTCAAGGCC
AATTTTGAAGAGGCTATATTTTCCAAGTATATAGGAAATGTCAACACACACGTGGATGAGTACATGCTGGAAGC
AGTGGACCACTACGCAGGCCAACTAGCCACCCTAGATATCAGCACTGAACCAATGAAACTGGAGGACGCAGTGT
ACGGTACCGAGGGTCTTGAGGCGCTTGATCTAACAACGAGTGCCGGTTACCCATATGTTGCACTGGGTATCAAG
AAGAGGGACATCCTCTCTAAGAAGACTAAGGACCTAACAAAGTTAAAGGAATGTATGGACAAGTATGGCCTGAA
CCTACCAATGGTGACTTATGTAAAAGATGAGCTCAGGTCCATAGAGAAGGTAGCGAAAGGAAAGTCTAGGCTGA
TTGAGGCGTCCAGTTTGAATGATTCAGTGGCGATGAGACAGACATTTGGTAATCTGTACAAAACTTTCCACCTA
AACCCAGGGGTTGTGACTGGTAGTGCTGTTGGGTGTGACCCAGACCTCTTTTGGAGCAAGATACCAGTGATGTT
AGATGGACATCTYATAGCATTTGATTACTCTGGGTACGATGCTAGCTTAAGCCCTGTCTGGTTTGCTTGCCTAA
AAATGTTACTTGAGAAGCTTGGATACACGCACAAAGAGACAAACTACATTGACTACTTGTGCAACTCCCATCAC
CTGTACAGGGATAAACATTACTTTGTGAGGGGTGGCATGCCCTCGGGATGTTCTGGTACCAGTATTTTCAACTC
AATGATTAACAATATCATAATTAGGACACTAATGCTAAAAGTGTACAAAGGGATTGACTTGGACCAATTCAGGA
TGATCGCATATGGTGATGATGTGATCGCATCGTACCCATGGCCTATAGATGCATCTTTACTCGCTGAAGCTGGT
AAGGGTTACGGGCTGATCATGACACCAGCAGATAAGGGAGAGTGCTTTAACGAAGTTACCTGGACCAACGTCAC
TTTCCTAAAGAGGTATTTTAGAGCAGATGAACAGTACCCCTTCCTGGTGCATCCTGTTATGCCCATGAAAGACA
TACACGAATCAATTAGATGGACCAAGGATCCAAAGAACACCCAAGATCACGTGCGCTCACTGTGTCTATTAGCT
TGGCATAACGGGGAGCACGAATATGAGGAGTTCATCCGTAAAATTAGAAGCGTCCCAGTCGGACGTTGTTTGAC
CCTCCCCGCGTTTTCAACTCTACGCAGGAAGTGGTTGGACTCCTTTTAGATTAGAGACAATTTGAAATAATTTA
GATTGGCTCAACCCTACTGTGCTAACCGAACCAGATAACGGTACAGTAGGGGTAAATTCTCCGCATTCGGTG

**>CVB3 amino acid translation, canonical reading frame**

MGAQVSTQKTGAHETGLNASGNSIIHYTNINYYKDAASNSANRQDFTQDPGKFTEPVKDIMIKSLPALNSPTVE
ECGYSDRARSITLGNSTITTQECANVVVGYGVWPDYLKDSEATAEDQPTQPDVATCRFYTLDSVQWQKTSPGWW
WKLPDALSNLGLFGQNMQYHYLGRTGYTVHVQCNASKFHQGCLLVVCVPEAEMGCATLDNTPSSAELLGGDSAK
EFADKPVASGSNKLVQRVVYNAGMGVGVGNLTIFPHQWINLRTNNSATIVMPYTNSVPMDNMFRHNNVTLMVIP
FVPLDYCPGSTTYVPITVTIAPMCAEYNGLRLAGHQGLPTMNTPGSCQFLTSDDFQSPSAMPQYDVTPEMRIPG
EVKNLMEIAEVDSVVPVQNVGEKVNSMEAYQIPVRSNEGSGTQVFGFPLQPGYSSVFSRTLLGEILNYYTHWSG
SIKLTFMFCGSAMATGKFLLAYSPPGAGAXTKRVDAMLGTHVIWDVGLQSSCVLCIPWISQTHYRYVASDEYTA
GGXITCWYQTNIVVPADAQSSCYIMCFVSACNDFSVRLLKDTPFISQENFFQGPVEDAITAAIGRVADTVGTGP
TNSEAIPALTAAETGHTSQVVPGDTMQTRHVKNYHSRSESTIENFLCRSACVYFTEYENSGAKRYAEWVLTPRQ
AAQLRRKLEFFTYVRFDLELTFVITSTQQPSTTQNQDAQILTHQIMYVPPGGPVPDKVDSYVWQTSTNPSVFWT
EGNAPPRMSIPFLSIGNAYSNFYDGWSEFSRNGVYGINTLNNMGTLYARHVNAGSTGPIKSTIRIYFKPKHVKA
WIPRPPRLCQYEKAKNVNFQPSGVTTTRQSITTMTNTGAFGQQSGAVYVXNYRVVNRXXATSADWQNCVWESYN
RDLLVSTTTAHGCDIIARCQCTTGVYFCASKNKHYPISFEGPXLVEVQESEYYPRRYQSHVLLAAGFSEPGDCG
GILRCEHGVIGIVTMGGEGVVGFADIRDLLWLEDDAMEQGVKDYVEQLGNAFGSGFTNQICEQVNLLKESLVGQ
DSILEKSLKALVKIISALVIVVRNHDDLITVTATLALIGCTSSPWRWLKQKXSQYYGIPMAERQNNSWLKKFTE
MTNACKGMEWIAVKIQKFIEWLKVKILPEVREKHEFLNRLKQLPLLESQIATIEQSAPSQSDQEQLFSNVQYFA
HYCRKYAPLYAAEAKRVFSLEKKMSNYIQFKSKCRIEPVCLLLHGSPGAGKSVATNLIGRSLAEKLNSSVYSLP
PDPDHFDGYKQQAVVIMDDLCQNPDGKDVSLFCQMVSSVDFVPPMAALEEKGILFTSPFVLASTNAGSINAPTV
SDSRALARRFHFDMNIEVISMYSQNGKINMPMSVKTCDDECCPVNFKKCCPLVCGKAIQFIDRRTQVRYSLDML
VTEMFREYNHRHSVGTTLEALFQGPPVYREIKISVAPETPPPPAIADLLKSVDSEAVREYCKEKGWLVPEINST
LQIEKHVSRAFICLQALTTFVSVAGIIYIIYKLFAGFQXAYTGVPNQKPRVPTLRQAKVQGPAFEFAVAMMKRN
SSTVKTEYGEFTMLGIYDRWAVLPRHAKPGPTILMNDQEVGVLDAKELVDKDGTNLELTLLKLNRNEKFRDIRG
FLAKEEVEVNEAVLAINTSKFPNMYIPVGQVTEYGFLNLGGTPTKRMLMYNFPTRAGQCGGVLMSTGKVLGIHV
GGNGHQGFSAALLKHYFNDEQGEIEFIESSKDAGFPVINTPSKTKLEPSVFHQVFEGNKEPAVLRSGDPRLKAN
FEEAIFSKYIGNVNTHVDEYMLEAVDHYAGQLATLDISTEPMKLEDAVYGTEGLEALDLTTSAGYPYVALGIKK
RDILSKKTKDLTKLKECMDKYGLNLPMVTYVKDELRSIEKVAKGKSRLIEASSLNDSVAMRQTFGNLYKTFHLN
PGVVTGSAVGCDPDLFWSKIPVMLDGHXIAFDYSGYDASLSPVWFACLKMLLEKLGYTHKETNYIDYLCNSHHL
YRDKHYFVRGGMPSGCSGTSIFNSMINNIIIRTLMLKVYKGIDLDQFRMIAYGDDVIASYPWPIDASLLAEAGK
GYGLIMTPADKGECFNEVTWTNVTFLKRYFRADEQYPFLVHPVMPMKDIHESIRWTKDPKNTQDHVRSLCLLAW
HNGEHEYEEFIRKIRSVPVGRCLTLPAFSTLRRKWLDSF
